# Three-dimensional residual channel attention networks denoise and sharpen fluorescence microscopy image volumes

**DOI:** 10.1101/2020.08.27.270439

**Authors:** Jiji Chen, Hideki Sasaki, Hoyin Lai, Yijun Su, Jiamin Liu, Yicong Wu, Alexander Zhovmer, Christian A. Combs, Ivan Rey-Suarez, Hungyu Chang, Chi Chou Huang, Xuesong Li, Min Guo, Srineil Nizambad, Arpita Upadhyaya, Shih-Jong J. Lee, Luciano A.G. Lucas, Hari Shroff

## Abstract

We demonstrate residual channel attention networks (RCAN) for restoring and enhancing volumetric time-lapse (4D) fluorescence microscopy data. First, we modify RCAN to handle image volumes, showing that our network enables denoising competitive with three other state-of-the-art neural networks. We use RCAN to restore noisy 4D super-resolution data, enabling image capture over tens of thousands of images (thousands of volumes) without apparent photobleaching. Second, using simulations we show that RCAN enables class-leading resolution enhancement, superior to other networks. Third, we exploit RCAN for denoising and resolution improvement in confocal microscopy, enabling ∼2.5-fold lateral resolution enhancement using stimulated emission depletion (STED) microscopy ground truth. Fourth, we develop methods to improve spatial resolution in structured illumination microscopy using expansion microscopy ground truth, achieving improvements of ∼1.4-fold laterally and ∼3.4-fold axially. Finally, we characterize the limits of denoising and resolution enhancement, suggesting practical benchmarks for evaluating and further enhancing network performance.

## Introduction

All fluorescence microscopes suffer drawbacks and tradeoffs because they partition a finite signal budget in space and time. These limitations manifest when comparing different microscope types (e.g., three-dimensional structured illumination microscopy^1^ (SIM) offers better spatial resolution than high numerical aperture light sheet microscopy^2^, but worse photobleaching); different implementations of the same microscope type (e.g., traditional implementations of SIM offer better spatial resolution than instant SIM (iSIM)^3^, but worse depth penetration and lower speed^4^); and within the same microscope (longer exposures and bigger pixels increase signal-to-noise ratio (SNR) at the expense of speed and resolution^5^). Performance tradeoffs are especially severe^6^ when considering live-cell super-resolution microscopy applications, in which the desired spatiotemporal resolution must be balanced against sample health^7^.

Deep learning^8^, which harnesses neural networks for data-driven statistical inference, has emerged as a promising method for alleviating drawbacks in fluorescence microscopy. Content-aware image restoration (CARE^9^) networks use the popular U-net^10^ neural network architecture in conjunction with synthetic, semi-synthetic and physically acquired training data to improve resolution, resolution isotropy, and signal-to-noise ratio in fluorescence images. U-nets have also been incorporated into generative adversarial networks (GAN^11^) that enable cross-modality super-resolution microscopy, transforming confocal images into STED images^12^ or transforming a series of widefield or sparse localization microscopy images into high resolution localization microscopy images^13^. Other recent examples include denoising confocal^14^ or SIM^15^ data and deconvolving light-sheet data^16^.

Here we investigate the use of an alternative network architecture, the residual channel attention network (RCAN)^17^, for use in super-resolution microscopy applications. RCAN has been shown to preferentially learn high spatial frequency detail within natural scene images, but this capability has not been exploited for image restoration in fluorescence microscopy applications, nor on longitudinally acquired image volumes. First, we modify RCAN for 3D applications, showing that it matches or exceeds the performance of previous networks in denoising fluorescence microscopy data. We apply this capability for super-resolution imaging over thousands of image volumes (tens of thousands of images). Second, we characterize RCAN and other networks in terms of their ability to extend resolution, finding that RCAN provides better resolution enhancement than alternatives, especially along the axial dimension. Finally, we demonstrate 4-5 fold volumetric resolution improvement in multiple fixed- and live-cell samples when using stimulated emission depletion (STED)- and expansion^18^-microscopy ground truth to train RCAN models.

## Results

### RCAN enables super-resolution imaging over thousands of volumes

The original RCAN was proposed specifically for resolution enhancement^17^. A key challenge in this task is the need to bypass abundant low-resolution information in the input image in favor of high-resolution prediction. The RCAN architecture achieves this by employing multiple skip connections between network layers to bypass low-resolution content, as well as a ‘channel-attention’ mechanism^19^ that emphasizes the more relevant feature channels, preventing low resolution features from dominating the prediction. We modified the original RCAN architecture to handle image volumes rather than images, also improving network efficiency so that our modified 3D RCAN model fits within graphics processing unit (GPU) memory (**Fig. 1a, Methods, Supplementary Note 1**).

**Fig. 1.**
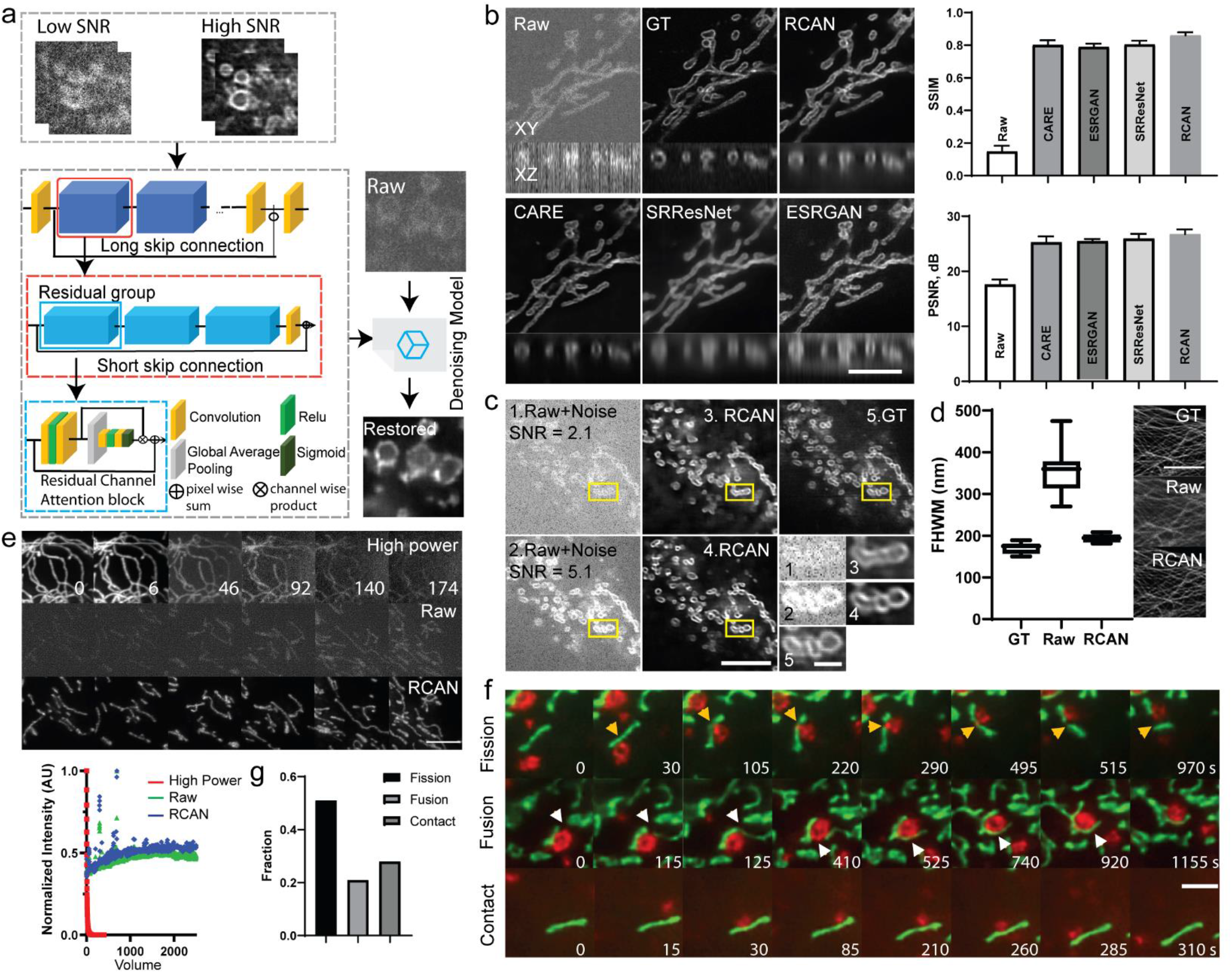
Residual channel attention networks denoise super-resolution data. **a)** Residual channel attention network (RCAN) architecture used throughout this work. Matched low and high SNR image volumes are used to train our RCAN, a residual in residual structure which consists of several residual groups (dark blue, red outline) with long skip connections. Each residual group itself contains additional residual channel attention blocks (RCAB, light blue, blue outline) with short skip connections, convolution, rectified linear unit (ReLu), sigmoid, and pooling operations. Long and short skip connections, as well as short-cuts within the residual blocks, allow abundant low-frequency information to be bypassed through such identity-based skip connections, facilitating the learning of high frequency information. A channel attention mechanism within the RCAB further aids the representational ability of the network in learning high-resolution information. **b)** *Left*: noisy raw instant SIM (iSIM) data acquired with low-intensity illumination, low-noise deconvolved ground truth (GT) data acquired with high-intensity illumination, RCAN, CARE, SRResNet, and ESRGAN output. Lateral (upper) and axial (lower) cross sections are shown. Samples are fixed U2OS cells expressing mEmerald-Tomm20, imaged via iSIM. *Right*: Comparison of network output using structural similarity index (SSIM) and peak signal-to-noise-ratio (PSNR). Means and standard deviations are reported, obtained from *N* = 10 planes from one volume. See also **Supplementary Figs. 1, 2. c**) RCAN performance at different input SNR levels, simulated by adding Gaussian and Poisson noise to raw input. Noisy raw input data at SNR 2.1 (top row) and 5.1 (bottom row) were used to generate predictions, which were then compared to ground truth. SNR values are calculated as mean of values within the yellow rectangular regions. Higher magnification views of mitochondria marked in yellow rectangular regions are shown at lower right. See also **Supplementary Fig. 5. d)** Full width at half maximum values (mean +/− standard deviations) from 10 microtubule filaments for deconvolved, high SNR ground truth (GT); noisy iSIM input (‘Raw’); and network output (‘RCAN’). **e**) RCAN denoising enables collection of thousands of iSIM volumes without photobleaching. Mitochondria in live U2OS cells were labeled with pShooter pEF-Myc-mito-GFP and imaged with high (360 W/cm^2^) and low (4.2 W/cm^2^) intensity illumination. Top row: selected examples at high illumination power, illustrating severe photobleaching. Middle row: selected examples from a different cell, imaged at low illumination power, illustrating low SNR (‘Raw’). Bottom row: RCAN output given low SNR input. Numbers in top row indicate volume #. Graph quantifies normalized signal in each case, ‘jumps’ in Raw and RCAN signal correspond to manual refocusing during acquisition. Maximum intensity projections are shown. See also **Supplementary Videos 1, 2, Supplementary Fig. 6, 7. f**) Dual-color imaging of mitochondria (green, pShooter pEF-Myc-mito-GFP) and lysosomes (mApple-Lamp1) in live U2OS cells. RCAN output illustrating mitochondrial fission (orange arrowheads), mitochondrial fusion (white arrowheads), and mitochondrial-lysosomal contacts. Single lateral planes are shown. See also **Supplementary Video 3. g**) Graph shows quantification of fission, fusion, and contact events quantified from 16 cells. All scale bars: 5 μm, except 1 μm for higher magnification views shown in **c)**.

To investigate RCAN denoising performance on fluorescence data, we began by acquiring matched pairs of low- and high-SNR iSIM volumes of fixed U2OS cells transfected with mEmerald-Tomm20 (**Methods, Supplementary Table 1, 2**), labeling the outer mitochondrial membrane (**Fig. 1b**). We programmed our acousto-optic tunable filter to rapidly switch between low (4.2 W/cm^2^) and high (457 W/cm^2^) intensity illumination, rapidly acquiring 35 low SNR raw volumes and matching high SNR data, which we deconvolved to yield high SNR ‘ground truth’. We then used 30 of these volumes for training and held out 5 volumes for testing network performance. Using the same training and test data, we compared four networks: RCAN, CARE, SRResNET^20^, and ESRGAN^21^. SRResNet and ESRGAN are both class-leading deep residual networks used in image super-resolution, with ESRGAN winning the 2018 Perceptual Image Restoration and Manipulation challenge on perceptual image super-resolution^22^.

For the mEmerald-Tomm20 label, RCAN, CARE, ESRGAN, and SRResNET predictions all provided clear improvements in visual appearance, structural similarity index (SSIM) and peak signal-to-noise-ratio (PSNR) metrics relative to the raw input (**Fig. 1b**), also outperforming direct deconvolution on the noisy input data (**Supplementary Fig. 1**). The RCAN output provided PSNR and SSIM values competitive with the other networks (**Fig. 1b**), prompting us to investigate whether this performance held for other organelles. We thus conducted similar experiments for fixed U2OS cells with labeled actin, endoplasmic reticulum (ER), golgi, lysosomes, and microtubules (**Supplementary Fig. 2**), acquiring 15-23 volumes of training data and training independent networks for each organelle. In almost all cases, RCAN performance met or exceeded the other networks (**Supplementary Fig. 3, Supplementary Table 3**).

An essential consideration when using any deep learning method is understanding when network performance deteriorates. Independently training an ensemble of networks and computing measures of network disagreement can provide insight into this issue^9,16^, yet such measures were not generally predictive of disagreement between ground truth and RCAN output (**Supplementary Fig. 4**). Instead, we found that estimating the per-pixel SNR in the raw input (**Methods, Supplementary Fig. 4**) seemed to better correlate with network performance, with extremely noisy input generating a poor prediction, as intuitively expected. For example, for the mEmerald-Tomm20 and ERmoxGFP labels, we observed obvious artifacts when input SNR dropped below ∼3 (**Fig. 1c**). We observed similar effects when using synthetic spherical phantoms in the presence of large noise levels (**Supplementary Fig. 5**).

We also examined linearity and spatial resolution in the denoised RCAN predictions. We verified that the RCAN output reflected spatial variations in fluorescence intensity evident in the input data, demonstrating that linearity is preserved (**Supplementary Fig. 6**). To estimate spatial resolution, we examined the apparent full width at half maximum of 10 labeled microtubule filaments in noisy raw input; high SNR deconvolved ground truth; and the RCAN prediction (**Fig. 1d**). While lateral resolution was not recovered to the extent evident in the ground truth (170 +/− 13 nm, mean +/− standard deviation), predictions offered noticeable resolution improvement compared to the input data (194 +/− 9 nm RCAN vs. 353 +/− 58 nm input).

Next, we tested the performance of RCAN on live cells, for extended volumetric time-lapse (4D) imaging applications. At high SNR, relatively few volumes can be obtained with iSIM, due to significant volumetric bleaching. For example, when volumetrically imaging pShooter pEF-Myc-mito-GFP (labeling the mitochondrial matrix) in live U2OS cells every 5.6 s at high intensity (360 W/cm^2^, **Fig. 1e, Supplementary Video 1**), only seven volumes could be acquired before fluorescence dropped to half its initial value. Lowering the illumination intensity to 4.2 W/cm^2^ so that photobleaching is negligible compared to the rate of protein synthesis circumvents this problem, but the resulting low SNR usually renders the data unusable (**Fig. 1e**). To determine whether deep learning could help to address this tradeoff between SNR and imaging duration, we accumulated 36 matched low (4.2 W/cm^2^)/high intensity (457 W/cm^2^) volumes on fixed cells, and trained an RCAN model, which we then tested on our low SNR live data. This approach enabled super-resolution imaging over an extended duration, allowing capture of 2600 image volumes (∼50,000 images, 2.2 W/cm^2^) acquired every 5.6 s over four hours with no detectable photobleaching and an apparent increase in fluorescence signal over the course of the recording (**Fig. 1e, Supplementary Video 2**). The restored image quality was sufficiently high that individual mitochondria could be manually segmented, a task difficult or impossible on the raw input data (**Supplementary Fig. 7**). To our knowledge, light-sheet microscopy is the only technique capable of generating 4D data of similar quality and duration, but the sub-200 nm spatial resolution of our method is better than that of high-NA light-sheet microscopy^23^. In another application, a dual-color example, we applied the same strategy to imaging pShooter pEF-Myc-mito-GFP in conjunction with mApple-LAMP1 labeled lysosomes. In this case, we obtained ∼300 super-resolution volumes recorded every 5.1 s in a representative cell (**Supplementary Video 3**), allowing inspection (**Fig 1f**) of mitochondrial fission and fusion near lysosomal contacts. Manually quantifying these events from 16 cells, we found that fission occurred ∼2.5x as often as fusion (**Fig. 1g**).

### Estimating the resolution enhancement offered by deep learning

In addition to denoising fluorescence images, deep learning can also be used for resolution enhancement^9,12,13^. We were curious about the extent to which RCAN (and other networks) could retrieve resolution degraded by the optical system, since this capability has not been systematically investigated. We were particularly interested in understanding when network performance breaks down, i.e., how much blurring is too much. To empirically assess the relative performance of different networks, we simulated ground truth noiseless spherical phantoms and subjected them to increasing amounts of blur (**Fig. 2, Supplementary Videos 4-6**). We trained RCAN, CARE, SRResNet, and ESRGAN networks with the same 23 matched volumes of ground truth and blurred data, and then challenged each network with 7 volumes of previously unseen test data (**Fig. 2a-c, Supplementary Figure 8**).

**Fig. 2.**
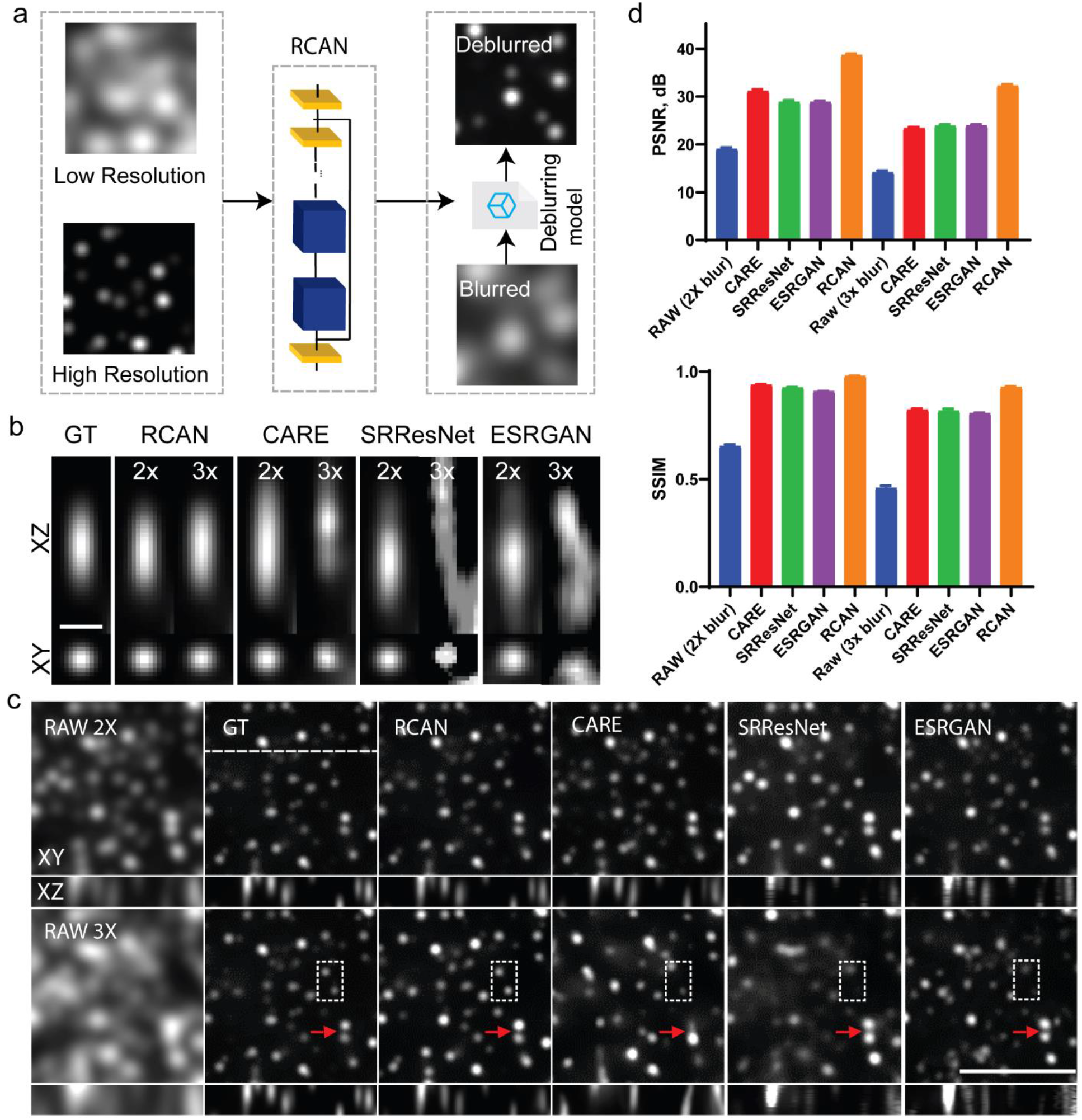
RCAN resolution enhancement assayed with simulated spherical phantoms. **a)** Noiseless mages of simulated spherical phantoms were created (High Resolution) and blurred (Low Resolution), generating matched volumes for RCAN training. Blurred volumes unseen by the trained network were then tested to evaluate deblurring performance. **b)** Examples of RCAN, CARE, SRResNet and ESRGAN performance on increasingly blurred data (blurred with a kernel 2x and 3x larger than the iSIM PSF used for ground truth (GT) data). Axial (top row) and lateral (bottom row) cross sections are shown. Networks are compared on the same test object, a sub-resolution sphere that approximates the iSIM PSF after blurring (GT, shown in leftmost column). Scale bar: 40 pixels. See also **Supplementary Fig. 8. c)** Additional examples of input data after progressively more severe blur (RAW, left column, with blurring kernels 2x and 3x the size of the iSIM PSF indicated in successive rows). Ground truth and different network outputs (right column) are also shown. Scale bar: 100 pixels, lateral (XY, top images) and axial slices (XZ, bottom images) along the dotted horizontal line are shown. Dotted rectangles and red arrows highlight features for comparison across the different networks. See also **Supplementary Fig. 9, Supplementary Videos 4-6. d)** SSIM (top) and PSNR (bottom) for data shown in **c)**. Means and standard deviations from 8 measurements are shown, see also **Supplementary Table 4**.

The RCAN generated plausible reconstructions even with blurring 3-fold greater (in all spatial dimensions) than the iSIM PSF (**Fig. 2b**), largely preserving the size of the smallest particles (**Fig. 2b,c**). However, RCAN performance degraded with increasingly blurry input, with SSIM and PSNR decaying from 0.98 to 0.93 and 38 dB to 32 dB for two- to three-fold blur, with other networks also showing worsened performance at increasing blur (**Fig. 2d, Supplementary Table 4**). Compared to the other networks, RCAN predictions offered improved resolution along the axial dimension (**Fig. 2b, c, Supplementary Fig. 8**), and superior SSIM and PSNR (**Fig. 2d, Supplementary Table 4**). We noticed obvious artifacts in all networks at 4x blur, suggesting an effective limit for deblurring with deep learning (**Supplementary Fig. 9, Supplementary Video 6**).

### Using RCAN for confocal to STED resolution enhancement in fixed and live cells

Since the noiseless spherical phantoms suggested that RCAN provides class-leading performance for resolution enhancement, we sought to benchmark RCAN performance using noisy experimental data. As a first test, we studied the ability to ‘transform’ confocal volumes into volumes with STED-like spatial resolution (**Fig. 3**), which is attractive because confocal imaging provides gentler, higher SNR imaging than STED microscopy but worse spatial resolution. Such ‘cross-modality’ super-resolution has been demonstrated before with GANs, but only with 2D images obtained from fixed cells^12^.

**Fig. 3.**
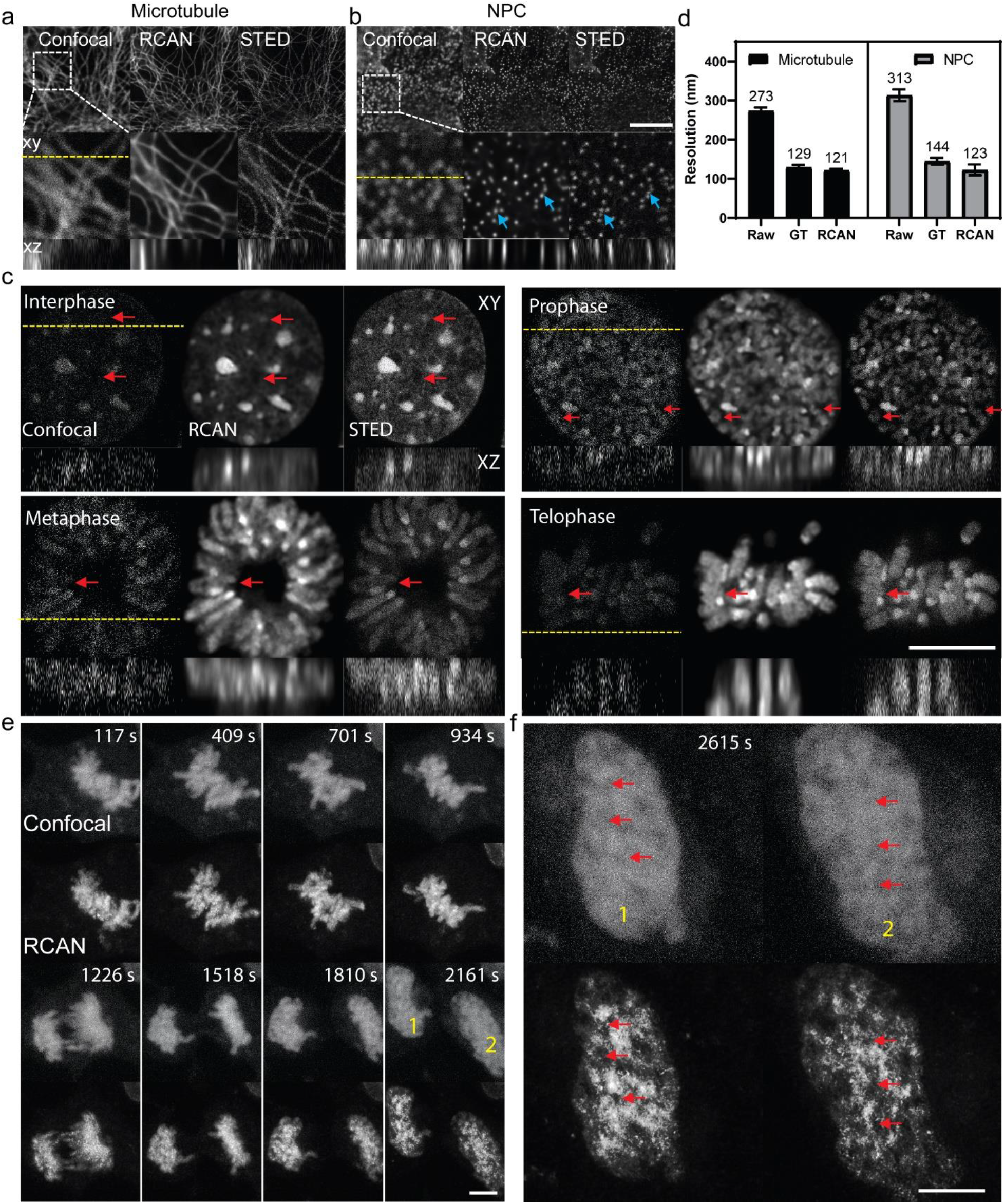
Confocal- to STED- microscopy restoration with RCAN. **a**) Example confocal input (left), RCAN prediction (middle) and ground truth STED (right) images for fixed mouse embryonic fibroblast (MEF) cells with microtubules stained with ATTO647-secondary antibodies against anti-α-tubulin primary antibodies (**a**), nuclear pore complexes (NPCs) stained with Alexa Fluor 594-secondary antibodies against anti-NPC primary antibodies (**b**) and nuclei stained with SiR-DNA (**c**). Higher magnification views of the dotted rectangular regions are shown below **a, b**, and axial reslices along yellow dotted lines marked in the lateral images are shown for **a-c**. Blue arrows in **b)** highlight areas of discrepancy between RCAN output and ground truth data while red arrows in **c)** are intended to highlight areas that are predicted well by RCAN but barely visible in the raw data. See also **Supplementary Fig. 10, 12, 13**. Phases of the cell cycle are also indicated in **c**). See also **Supplementary Fig. 14**. (**d**) Average image resolution in microtubule (left) and NPC (right) images obtained from decorrelation analysis. Means (also shown above each column) and standard deviations (from *N* = 18 image planes) are shown for raw confocal input, ground truth STED, and RCAN output. (**e**) Live MEF cells stained with SiR-DNA were imaged in resonant confocal mode (top) and the RCAN model trained on fixed datasets similar to those shown in (**c**) was applied to yield predictions (bottom). Single planes from volumetric time series are shown See also **Supplementary Video 7, 8. f)** Higher magnification view from series in **e)** 2615 s after the start of imaging, corresponding to nuclei marked 1, 2 in **e**). Red arrows highlight areas absent SiR-DNA signal that are more easily defined in RCAN prediction vs. confocal data. All scale bars: 5 μm.

We collected training data (22-26 volumes, **Supplementary Table 2**) on fixed, fluorescently labeled mouse embryonic fibroblast cells using a commercial Leica SP8 3X STED microscope (**Fig. 3a-c**). This system was particularly convenient as the STED images could be acquired immediately after the confocal images, on the same instrument. We imaged fixed mouse embryonic fibroblasts, immunostained with ATTO647-secondary antibodies against anti-α-tubulin primary antibodies for marking microtubules (**Fig. 3a**); and with Alexa Fluor 594-secondary antibodies against anti-NPC primary antibodies, marking nuclear pores (**Fig. 3b**). Next, we trained RCAN models and applied them to unseen data (**Supplementary Fig. 10**), using a modified decorrelation analysis^24^ (**Methods, Supplementary Fig. 11**) to estimate average spatial resolution. Confocal spatial resolution was 273 +/− 9 nm (*N* = 18 images used for these measurements) in the microtubule dataset and 313 +/− 14 nm in the pore dataset, with STED microscopy providing ∼2-fold improvement in resolution (129 +/− 6 nm for microtubules, 144 +/− 9 nm for the pores) and the RCAN prediction providing similar gains (121 +/− 4 nm microtubules, 123 +/− 14 nm nuclear pores, **Fig. 3d**) that could not be matched by deconvolving the confocal data (**Supplementary Fig. 12**). We suspect that the slight improvement in spatial resolution in RCAN output relative to the STED ground truth is because the RCAN denoised the data as well as improved resolution, resulting in higher SNR than the STED ground truth. Close examination of the RCAN prediction for nuclear pores revealed slight differences in pore placement relative to the STED microscopy ground truth. We suspect that this result is due to slight differences in image registration between the confocal and STED data (**Supplementary Fig. 13**), perhaps due to sample drift between acquisitions or slight instrument misalignment. Applying an affine registration between the confocal and STED training data improved agreement between the confocal and STED data, improving network output (**Supplementary Fig. 13**). However, small deviations in nuclear pore placement between the ground truth STED and RCAN predictions were still evident.

We also examined a third label, SiR-DNA, a DNA stain well suited for labeling live and fixed cells in both confocal and STED microscopy^25^. Collecting matched confocal and STED volumes on fixed nuclei in a variety of mitotic stages enabled us to train a robust RCAN model that produced predictions on different nuclear morphologies (**Fig. 3c, Supplementary Fig. 14**) that were sharper and less noisy than confocal input. Improvement relative to the confocal data was particularly striking in the axial dimension (**Fig. 3c**). Given the quality of these reconstructions, we wondered whether the same RCAN model could be adapted for transfer learning on live samples.

Point-scanning confocal imaging can produce time-lapse volumetric recordings of living cells at SNR much higher than STED microscopy, given that more signal is collected per pixel. Nevertheless, even confocal microscopy recordings are quite noisy if high speed acquisitions are acquired. To demonstrate that our RCAN model trained on fixed cells could simultaneously denoise and improve resolution in live cells, we acquired noisy resonant confocal recordings of dividing cells labeled with SiR-DNA (**Fig. 3e**). Our illumination conditions were sufficiently gentle and rapid that we could acquire tens of imaging volumes without obvious bleaching or motion blur (**Supplementary Video 7**). Although the raw resonant confocal data poorly defined nuclei and chromosomes, these structures were clearly resolved in the RCAN predictions (**Fig. 3e, Supplementary Video 7**). The RCAN also better captured chromosome decondensation and the return to interphase DNA structure (**Fig. 3f**, see also additional interphase cell comparisons in **Supplementary Video 8**).

### Using expansion microscopy to improve iSIM resolution in fixed and live cells

Our success in using fixed STED training data to improve the spatial resolution of confocal microscopy made us wonder whether a similar strategy could be used to improve spatial resolution in iSIM. Since our iSIM did not inherently possess a means to image specimens at higher resolution than that of the base microscope, we used expansion microscopy (ExM^18^) to provide higher-resolution training data (**Fig. 4a**). ExM physically expands fixed tissue using a hydrogel and can improve resolution near-isotropically up to a factor given by the gel expansion. We used ultrastructure expansion microscopy (U-ExM^26^, a variant of the original ExM protocol) to expand mitochondria (immunolabeled with Rabbit-α-Tomm20 primary and Donkey-α-Rabbit Biotin secondary antibodies and Alexa Fluor 488 Streptavidin) and microtubules (labeled with Mouse-α-Tubulin primary and Donkey-α-Mouse Biotin secondary antibodies and Alexa Fluor 488 Streptavidin) in fixed U2OS cells by 3.2- and 4-fold, respectively (**Methods, Supplementary Fig. 15**), also developing protocols to locate and image the same region before- and after ExM with iSIM (**Supplementary Fig. 16, Methods**).

**Fig. 4.**
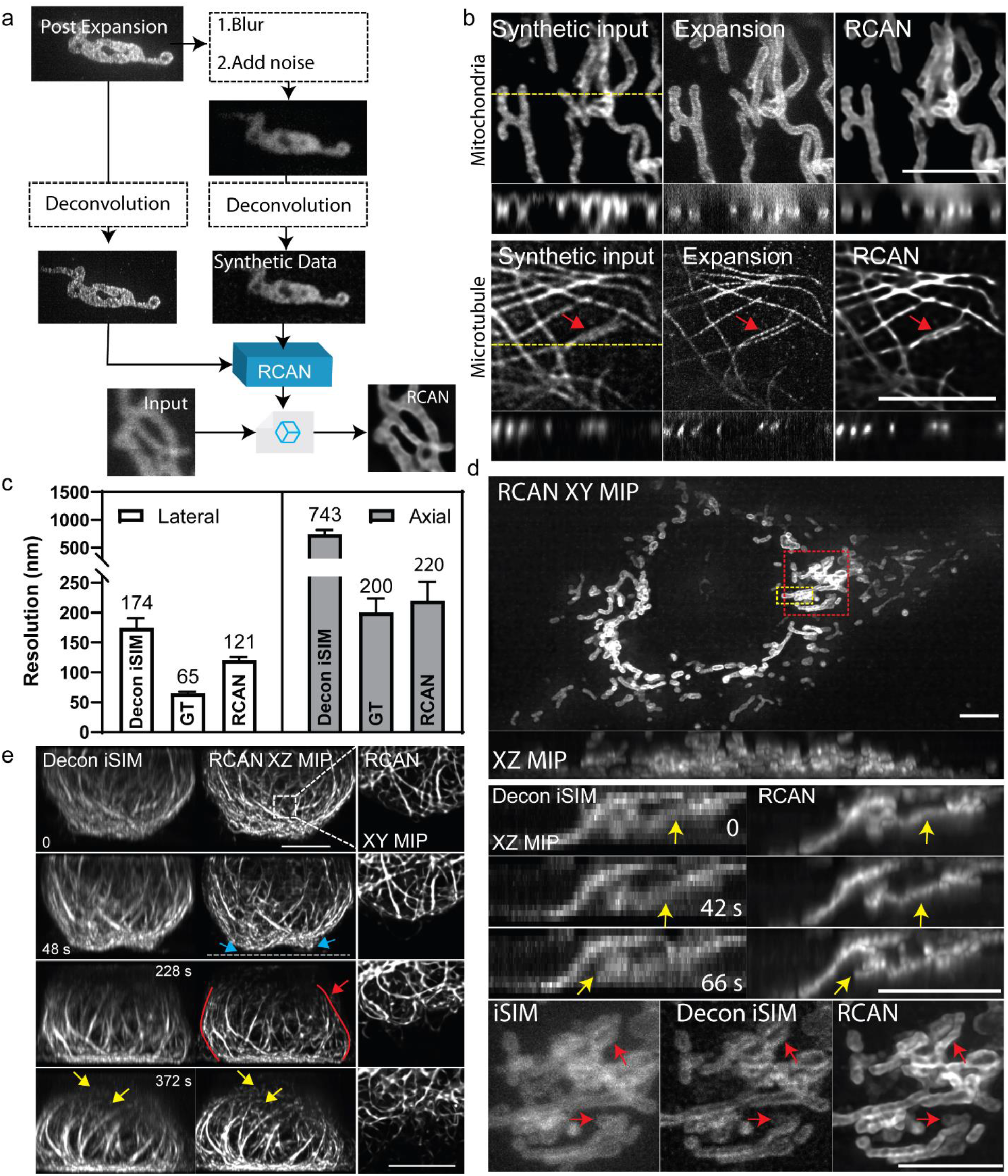
Using expansion microscopy to improve spatial resolution in fixed and live instant structured illumination microscopy (iSIM). **a)** Simplified schematic showing generation of synthetic data used for training RCAN network. Post-expansion data are acquired and deconvolved, generating ground truth data (left). Post-expansion data are also blurred, noise is added, and the resulting images are deconvolved to generate synthetic pre-expansion data (right). Ground truth and synthetic data are then used to train RCAN models for resolution enhancement on blurry input data (bottom). See also **Supplementary Fig. 19. b)** Example input data (either synthetic or experimental) not seen by network, mimicking deconvolved iSIM (left); expansion ground truth (middle); and RCAN predictions. Lateral and axial (taken along dotted line in lateral view) slices are shown for mitochondria (top, labeled with EGFP-Tomm20 in fixed, expanded U2OS cells) and microtubules (bottom, immunolabeled with Alexa Fluor 488 secondary against anti-α-tubulin primary antibody in fixed, expanded U2OS cells). See also **Supplementary Fig. 20** and **Supplementary Video 9. c)** Average resolution quantification from decorrelation analysis on microtubule samples. Lateral (left) and axial (right) values are shown for experimentally acquired deconvolved iSIM (left columns, 174 +/− 16 nm and 743 +/− 73 nm), ground truth expanded data (middle columns, 65 +/− 2 nm and 200 +/− 24 nm), and RCAN predictions (right columns, 120 +/− 5 nm and 220 +/− 31 nm). Note discontinuous representation of ordinate axis. Mean (shown also above each column) +/− standard deviations derived from *N* = 12 images are shown. See also **Supplementary Fig. 21. d**) Images from live U2OS cells expressing EGFP-Tomm20 were imaged with iSIM, deconvolved, and input into the trained RCAN model. Top: Overview lateral and axial maximum intensity projections (MIP) of first volume in time-series from RCAN prediction. Middle: higher magnification views of axial slice corresponding to yellow rectangular region in overview, comparing deconvolved iSIM input (left) and RCAN output. Yellow arrows highlight mitochondria that are better resolved with RCAN output than input data. Bottom: higher magnification views of red rectangular region in overview, comparing raw iSIM, deconvolved iSIM, and RCAN prediction. Red arrows highlight mitochondria better resolved with RCAN than iSIM. See also **Supplementary Videos 10, 11. e)** Images from live Jurkat T cells expressing EMTB-3XGFP were deconvolved and used as input to trained RCAN model. Left: selected axial MIPs at indicated time points, comparing deconvolved iSIM vs. RCAN output. Right: lateral MIPs, corresponding to dashed rectangular region in lefthand images. Blue arrowheads indicates deformation of lower cell cortex prior to T cell spreading, red arrow indicates approximate location of centrosome, red lines indicate asymmetric deformation of microtubule bundles surrounding the nucleus, and yellow arrows indicate microtubule filaments at the top of the cell better defined with RCAN vs iSIM. See also **Supplementary Videos 12, 13**. All scale bars: 5 μm.

We first attempted to directly register pre-ExM iSIM data to post-ExM data to build a training dataset suitable RCAN. Unfortunately, local distortions in the post-ExM data prevented the subpixel registration needed for accurate correspondence between pre- and post-ExM data, even when using landmark-based non-affine based registration methods (**Supplementary Fig. 17**). Instead, we digitally degraded the post-ExM data so that it resembled the lower resolution, pre-ExM iSIM data (**Fig. 4a**). Simply blurring the post-ExM data is insufficient, as blurring also oversmooths the background to the point that the images are noticeably smoother and less noisy than acquired pre-ExM iSIM data (**Supplementary Fig. 18**). Instead, we developed methods to match noise and background signal so that the digitally degraded post-ExM iSIM data better resembled deconvolved, pre-ExM iSIM data (**Supplementary Fig. 19, Methods**). This approach allowed us to register image pairs perfectly and to train RCAN models for microtubule and mitochondrial labels (**Methods, Supplementary Video 9, Supplementary Fig. 20**).

On fixed samples, the trained networks provided modest lateral resolution enhancement on synthetic data derived from ground truth images of expanded immunostained mitochondria and microtubules from fixed U2OS cells (**Fig. 4b**), allowing us to occasionally resolve closely spaced filaments otherwise blurred in the synthetic images (red arrows, **Fig. 4b**). However, the axial resolution enhancement offered by RCAN was more dramatic, showing clear improvement similar to the ground truth images. Using decorrelation analysis to estimate the degree of resolution enhancement on the microtubule data, we found that RCAN offered 1.5-fold increase laterally and 2.8-fold increase axially relative to the synthetic deconvolved data, compared to 2.2-fold improvement (lateral) and 3.5-fold (axial) offered by the ground truth data (**Supplementary Fig. 21**). We observed similar enhancements on experimentally acquired pre-expansion data: 1.4- and 3.4-fold improvement laterally and axially by the RCAN, versus 2.7-fold and 3.7-fold improvement in the ground truth data (**Fig. 4c**).

The improvements in fixed cells prompted us to apply our ExM-trained RCAN models to living cells imaged with iSIM in volumetric time-lapse sequences (**Fig. 4d, e, Supplementary Videos 10-13**). In a first example, we applied the RCAN to mitochondria labeled with EGFP-Tomm20 in live U2OS cells (**Fig. 4d, Supplementary Video 10**). Modest improvements in lateral resolution and contrast with RCAN offered better definition of individual mitochondria, including the void regions contained within the outer-mitochondrial space (**Fig. 4d**, red arrows). As with the fixed cells, improvements in axial views of the specimen were more dramatic (**Supplementary Video 11**), allowing us to discern closely packed mitochondria that were otherwise blurred in the deconvolved iSIM data (**Fig. 4d**, yellow arrows).

In a second, transfer-learning example, we applied our expansion-RCAN model derived from immunostained U2OS cells to live Jurkat T cells transiently expressing EMTB-3xGFP^27^, a protein that labels microtubule filaments. Jurkat T cells settled onto anti-CD3 coated activating coverslips (**Fig. 4e, Supplementary Videos 12-14**), which mimic antigen presenting cells and enable investigation of the early stages of immune synapse formation^28^. Dynamics and organization of the actin and microtubule cytoskeleton during cell spreading are important regulators of this phenomenon. The RCAN output offered clear views of the microtubule cytoskeleton during the initial stages of this dynamic process, including the deformation of microtubule bundles surrounding the nucleus. We observed pronounced deformation of the central microtubule bundles at the dorsal cell surface as spreading initiated (blue arrowheads), suggesting that these bundles may be anchored to the actin cortex. Anchoring of microtubules to the actin cortex allows the repositioning of the centrosome, a hallmark of immune synapse maturation^29^. Interestingly, we observed a higher deformation of the microtubule bundles on the right side of the cell shown in **Fig. 4e**, likely due to the forces that push and pull the centrosome towards the substrate (initially also located on the right side of the cell, red arrow at 228 s). RCAN output offered views with better resolution and contrast than the deconvolved iSIM input, particularly axially and towards the top of the cell. In some cases, dim or axially blurred filaments barely discerned in the input data were clearly resolved in the RCAN view (yellow arrows in **Fig. 4e, Supplementary Video 12, 14**).

## Discussion

Here we focused on 4D imaging applications, because sustained volumetric imaging over extended duration at diffraction-limited or better spatial resolution remains a major challenge in fluorescence microscopy. We have shown that RCAN denoises and deconvolves fluorescence microscopy image volumes with performance competitive to state-of-the-art neural networks (**Fig. 1**). In live 4D super-resolution applications, which typically exhibit pronounced bleaching that limits experiment duration, RCAN restoration allows the illumination to be turned down to a level where the rate of photobleaching is drastically reduced or even negligible. Unacceptably noisy images can be restored, allowing for extended volumetric imaging similar to that attained with light-sheet microscopy, but with better spatial resolution. We suspect that RCAN carefully combined with high-resolution, but noisy, confocal microscopy may thus challenge the current primacy of light-sheet microscopy, particularly when imaging thin samples. At the same time, we expect RCAN denoising to synergize with light-sheet microscopy, allowing even greater gains in experiment duration (or speed) with that technique. RCAN also deblurs images, with better performance than the other networks we’ve tested (**Fig. 2**). We used this feature to improve spatial resolution in confocal microscopy (**Fig. 3**), achieving 2.5-fold improvement in lateral resolution and iSIM data (**Fig. 4**), achieving 1.4-fold improvement laterally and ∼3-fold improvement axially.

Our findings highlight limitations of current neural networks and workflows and point the way to further improvements. First, on denoising applications we found ‘breaking points’ of the RCAN network at low input SNR. Estimating such input SNR may be useful in addition to computing measures of network disagreement^9^, especially given that the latter were not especially predictive of differences between ground truth and denoised data (**Supplementary Fig. 4**). Second, for resolution enhancement applications, our simulations on noiseless data revealed that all networks suffer noticeable deterioration when attempting to deblur at blur levels greater than 2-fold. Perhaps this explains why attempts to restore blurry microscopy images with neural networks have enabled only relatively modest levels of deblurring^9,14^. The fact that RCAN yielded better reconstructions than other networks even at 3-fold blurring suggests that network architecture itself may have substantial impact on deblurring performance. Our simulations also show that increased degradation in network output correlates with increased blur (**Fig. 2d**), implying caution is prudent when attempting extreme levels of deblurring. Exploring the fundamental limits of deblurring with neural networks would be an interesting avenue of further research. Third, practical factors still limit the performance of network output, suggesting that further improvement is possible. For the confocal-to-STED restorations, local deviations in spatial alignment between the training data pairs likely contribute to error in nuclear pore placement (**Supplementary Fig. 13**), suggesting that a local registration step during training would boost the quality of the restorations. For the expansion microscopy data, although we bypassed the need to finely register input and ground truth data by simulating pre-expansion data, improved registration schemes may enable direct use of experimentally derived pre- and post-expansion pairs. We suspect this would further improve the degree of resolution enhancement as complex noise and background variations in the data could be incorporated into the training procedure. We also expect that increasing label density would further improve the quality of our training data, as at the ∼65 nm resolution we achieved in the ground truth expansion data, stochastic variations in labelling were evident (**Supplementary Fig. 22**) and likely contribute an additional source of noise. Such improvements would probably also increase the SSIM and PSNR in the expansion predictions (**Supplementary Fig. 20**), which were markedly lower than in the confocal to STED predictions (**Supplementary Fig. 10**). Finally, achieving better spatial resolution in live samples usually demands corresponding improvements in temporal resolution, lest motion blur defeat gains in spatial resolution. We did not attempt to further increase the speed of our live recordings to account for this effect but doing so may result in sharper images.

Despite these caveats, the RCAN in its current form improves noisy super-resolution acquisitions, enabling image capture over tens of thousands of images; quantification, segmentation, and tracking of organelles and organelle dynamics; and prediction and inspection of fine details in confocal and iSIM data otherwise hidden by blur. We hope that our work inspires further advances in the rapidly developing field of image restoration.

## Author Contributions

Conceived project: S-J.J.L., L.A.G.L., H.Shroff. Designed experiments: J.C., Y.S., A.Z., C.C., I R-S., A.U., H.Shroff. Performed experiments: J.C., Y.S., A.Z., C.C., I.R-S. Developed and tested 3D RCAN: H.Sasaki, H.L., H.C., C.C.H., S-J J.L., L.A.G.L. Adapted and tested CARE, SRResNet, ESRGAN: J.C. and J.L. Wrote software for hardware control: X.L. Conceived, developed, and tested expansion microscopy pipeline: J.C., H.Sasaki, H.L., Y.S., J.L., Y.W., X.L., M.G., H.Shroff. Performed and analyzed simulations: J.C., J.L., Y.W., M.G., S.N., H.S. All authors analyzed data. Wrote paper: H.Shroff with input from all authors. Supervised research: A.U., S-J.J.L., L.A.G.L., H.Shroff.

## Acknowledgements

J.C., Y.W., and H.Shroff thank the Marine Biological Laboratory at Woods Hole for providing the Deep Learning for Microscopy Image Analysis Course as well as Florian Jug, Jenny Folkesson, and Patrick La Riviere for providing superb instruction. We thank William Bement for the gift of the EMTB-3XEGFP plasmid (also available as Addgene plasmid #26741), Justin Taraska for the LAMP1-GFP plasmid, George Patterson for the GalT-GFP plasmid, and Panagiotis Chandris for the pShooter pEF-Myc-mito-GFP plasmid. We also thank Patrick La Riviere and Hank Eden for their careful read and comments on this work. This research was supported by the intramural research programs of the National Institute of Biomedical Imaging and Bioengineering and the National Institute of Heart, Lung, and Blood within the National Institutes of Health and by an SBIR cooperative agreement of the National Institute of General Medical Sciences: 1U44GM136091-01. A.U. and I.R. acknowledge support from NIH R01 GM131054 and NSF PHY 1607645 grants.

## Disclaimer

The NIH, its staff, and officers do not recommend or endorse any company, product, or service.

## Methods

### Neural networks used for image restoration

#### 3D RCAN

The RCAN consists of multiple residual groups which themselves contain residual structure. Such ‘residual in residual’ structure forms a very deep network consisting of multiple residual groups with long skip connections (**Fig. 1a**). Each residual group also contains residual channel attention blocks (RCAB) with short skip connections. The long and short skip connections, as well as shortcuts within the residual blocks, allow low spatial frequency information to be bypassed, facilitating the prediction of high spatial frequency information. Additionally, a channel attention mechanism^19^ within the RCAB is used to adaptively rescale channel-wise features by considering interdependencies among channels, further improving the capability of the network to achieve higher resolution.

We extended the original RCAN^17^ to handle image volumes. Since 3D models with a large patch size may consume prohibitive GPU memory, we also changed various network parameters to ensure that our modified RCAN fits within GPU memory. These changes relative to the original RCAN model include: (1) we set the number of residual groups (RG) to G = 5 in the RIR structure; (2) in each RG, the RCAB number is set to 3; (3) the number of convolutional (Conv) layers in the shallow feature extraction and RIR structure is C = 32; (4) the Conv layer in channel-downscaling has C/r = 4 filters, where the reduction ratio r is set to 8; (5) all 2D Conv layers are replaced with 3D conv layers; (6) the upscaling module at the end of the network is omitted because network input and output have the same size in our case. In the original RCAN paper^17^, a small patch with size 48×48 is used for training. By contrast, we used a much larger patch size (256×256×16). We tried using a smaller patch size, but the training process was unstable and the results were poor. We suspect this is because microscopy images may show less high spatial frequency content than natural images, so a larger patch is necessary to extract enough gradient information for back-propagation.

The percentile-based image normalization proposed in the CARE manuscript^9^ is applied as a pre-processing step prior to training. In microscopy images, foreground objects of interest may be distributed sparsely. In such cases the model may overfit the background, failing to learn the structure of foreground objects if the entire image is used indiscriminately for training. To avoid overfitting, patches of the background were automatically rejected in favor of foreground patches during training. Background patch rejection is performed on the fly during data augmentation. We implemented training in a 3D version of RCAN using Keras^30^ with a TensorFlow^31^ backend. Each model was trained on two NVIDIA GeForce GTX 1080 Ti GPUs for 400 epochs, which took 1 day. Applying the denoising model on a 1920 x 1550 x 12 dataset using a desktop with a single GTX 1080 Ti GPU took ∼63.3 s per volume. This time also includes the time it takes to save the volume (with 32-bit output). On similar datasets with the same XY dimensions (but different number of Z-slices), applying the model took ∼3.9 s - 5.2 s per Z-slice. Further details are provided in **Supplementary Note 1** and **Supplementary Software**.

#### SRResNet and ESRGAN

SRResNet is a deep residual network for image super-resolution, which obtained state-of-the-art results in 2017^20^. Building on ResNet^32^, the SRResNet has 16 Residual Blocks (RB) with identical layout. Within each RB, there are two convolutional layers with small 3×3 kernels and 64 feature maps, followed by batch-normalization layers and a parametric rectified linear unit (ReLU) as activation function.

Generative adversarial networks^11^ (GAN) provide a powerful framework for generating plausible-looking natural images with high perceptual quality in computer vision applications. GANs are used in image super-resolution applications to favor solutions that resemble natural images^20^. Among such methods, enhanced super-resolution generative adversarial networks (ESRGAN^21^) won the first place in the Perceptual Image Restoration and Manipulation (PIRM) challenge on perceptual super-resolution in 2018^22^. Thus, we selected ESRGAN as an additional reference method to evaluate performance on fluorescence microcopy images.

The key concept underlying ESRGAN is to train a Generator *G* with the goal of fooling a Discriminator *D* that is trained to distinguish predicted high-resolution images from real high-resolution images. The Generator network *G* has 16 Residual in Residual Dense Blocks^21^ (RRDB) with identical layouts, which improves the RB design in SRResNet. RRDB has a residual-in-residual structure, where multi-level residual learning is used. In addition, RRDB contains dense blocks^33^, which increase network capacity due to the dense connections contained within each dense block.

The Discriminator network *D* is based on Relativistic GAN^34^. It has 8 convolutional layers with small 3×3 kernels as in the VGG network^35^ and the resulting feature maps are followed by two dense layers. A Relativistic average Discriminator^20^ (RaD) is used as the final activation function to predict the probability that a real high-resolution image is relatively more realistic than a fake high-resolution image.

In this work, we used the published SRResNet and ESRGAN (PyTorch implementation, https://github.com/xinntao/BasicSR) to process image volumes in a slice-by-slice manner. Before training, we normalized low-resolution (LR) and high-resolution (HR) images by percentile-based image normalization^9^ to reduce the effect of hot and dead pixels in the camera. Then we linearly rescaled the range of LR and HR images to [0,1]. SRResNet and ESRGAN networks were trained on an NVIDIA Quadro P6000 GPU. In all experiments (except the spherical phantoms), for each mini-batch, we cropped 16 random 480×480 overlapping image patches for training. Patches of background were not used for training. To determine whether a patch pair was from the background, we simply compared the mean intensity of the patch versus the whole image. If the mean intensity of the patch was less than 20% of the mean intensity of the whole image, the patch pair was not used for training. In spherical phantom experiments, we selected 16 random 2D image slices (256×256) for each mini-batch. For SRResNet, Adam optimization were used for all experiments with β_1_ = 0.9, β_2_ = 0.99, a learning rate of 2×10^−4^, and 10^5^ update iterations. During testing, batch-normalization update was turned off to obtain an output HR image that depended only on the input LR image. For ESRGAN, we used Adam optimization for all experiments with β_1_ = 0.9, β_2_ = 0.99. The Generator *G* and Discriminator *D* were alternately updated with learning rate initialized as 10^−4^ and decayed by a factor of 2 every 10^4^ updates. Training time was ∼8 hours for SRResNet and ∼12 hours for ESRGAN. Application usually took ∼60 s (SRResNet) to 120 s (ESRGAN) for the image volumes shown here.

#### CARE

The content aware restoration (CARE) framework has been described in detail.^9^ We implemented CARE through Keras and TensorFlow via GitHub (https://github.com/CSBDeep/CSBDeep). CARE networks were trained on an NVIDIA Titan RTX GPU card in a local workstation. Typically for each image volume, 2048 patches of size 128×128×8 were randomly cropped and used to train a CARE network with a learning rate of 2×10^−4^. From the extracted patches, 10% were used as validation data. The number of epochs for training is 200 and the mean absolute error (mae) was used as loss function. Training time for a given model was 8-12 hours, application of the model on a 1920×1550×28 sized image volume took ∼90 s.

For all networks, we evaluated the peak-signal-to-noise-ratio (PSNR) and the structural similarity index^29^ (SSIM) on normalized input, network output, and ground truth with built-in MATLAB (Mathworks) functions.

### Instant structured illumination microscopy (iSIM)

#### U2OS Cell Culture and transfection

U2OS cells were cultured and maintained at 37 C and 5% CO_2_ on glass bottom dishes (MatTek, P35G-1.5-14-C) in 1 mL of DMEM medium (Lonza, 12-604F) containing 10% FBS. At 40-60% confluency, cells were transfected with 100 μL of 1X PBS containing 2 μL of X-tremeGENE HP DNA transfection reagent (Sigma, 6366244001) and 2 μL plasmid DNA (300-400 ng/μL, see **Supplementary Table 1** for plasmid information) and maintained at 37C, 5 % CO_2_ for 1-2 days.

#### Immunofluorescence labeling

U2OS cells were fixed with 4% paraformaldehyde (Electron Microscopy Sciences, 15710) and 0.25% Glutaraldehyde (Sigma, G5882) in 1X PBS at room temperature (RT) for 15 minutes. Cells were rinsed 3 times with 1X PBS, and permeabilized by 0.1% Triton X-100 (Sigma, 93443) in 1X PBS for 1 minute. Cells were treated with 300μL Image-iT FX Signal enhancer (Thermofisher, R37107) for 30 minutes at RT followed by 30-minute blocking with 1% BSA/PBS (Thermofisher, 37525) at RT. Cells were then labeled with fluorescent antibodies and/or fluorescent streptavidin (see **Supplementary Table 1**) in 0.1% Triton X-100/PBS for 1 hour at RT. After antibody labeling, cells were washed 3 times with 0.1% Triton X-100 and stained by 4’, 6-diamidino-2-phenylindole (DAPI, Sigma, D9542) (1μg/mL) in 1X PBS for 5 minutes at RT. DAPI stain was used for expansion factor estimation and rapid cell or region localization throughout the Expansion Microscopy (ExM) process.

#### iSIM imaging for denoising

iSIM data was obtained on our previously reported home-built system^3^. A 60x NA 1.42 oil objective (Olympus) was used for all imaging, except the training data acquired for the iSIM to expansion microscopy cross-modality experiments (which used a 1.2 NA water immersion lens, described below in more detail). To obtain high and low SNR image pairs for training, high (usually 33 mW for 488 nm, 72 mW for 561 nm) and low powers (0.3 mW for 488 nm, 0.6 mW for 561 nm) were rapidly switched via an AOTF. Green and red fluorescence images were acquired with a filter wheel (Sutter, FG-LB10-BIQ and FG-LB10-NW) and notch filters (Semrock, NF03-488E-25 and NF03-561E-25). Samples were deposited on 35-mm-diameter high-precision 1.5 dish (Matek; P35G-0.170-14-C). For live cell imaging, the dishes were mounted within an incubation chamber (Okolab; H301-MINI) to maintain temperature at 37°C.

#### Estimating illumination intensity

A power meter (Thorlabs, PM100D) was used to measure the excitation laser power immediately prior to the objective. The average intensity was calculated using the measured intensity divided by the field of view (FOV, 106 μm by 68 μm).

#### Jurkat T Cell Culture, substrate preparation, and iSIM imaging

E6-1 Wild Type Jurkat cells were cultured in RPMI 1640 supplemented with 10% fetal bovine serum and 1% Penn-Strep antibiotics. Cells were transiently transfected with EMTB-3XGFP plasmid using the Neon (ThermoFisher Scientific) electroporation system two days before imaging, using the manufacturer’s protocol.

Coverslips attached to 8 well Labtek chambers were incubated in Poly-L-Lysine (PLL) at 0.01% W/V (Sigma Aldrich, St. Louis, MO) for 10 min. PLL was aspirated and the slide was left to dry for 1 hour at 37 °C. T cell activating antibody coating is performed by incubating the slides in a 10 μg/ml solution of anti-CD3 antibody (Hit-3a, eBiosciences, San Diego, CA) for 2 hours at 37 °C or overnight at 4 °C. Excess anti-CD3 was removed by washing with L-15 imaging media immediately prior to the experiment.

Imaging of live EMTB-3XGFP expressing Jurkat cells was performed at 37 °C using iSIM, with a 1.42 numerical aperture 60× lens (Olympus) and 488 nm laser for excitation using the same home-built system as above^3^. For volumetric live imaging, the exposure was set to 100 ms per slice, the spacing between slices to 250 nm, and the inter-volume temporal spacing to 12.3 s.

#### Linearity estimate

Linearity was assessed by measuring the intensity in different regions in maximum intensity projections (MIP) of raw images of fixed cells expressing U2OS cells expressing the mEmerald-Tomm20 label, and the corresponding RCAN predictions (**Supplementary Fig. 6**). Small regions of interest (ROIs, 8 by 8 pixels) were selected and the average intensity value in each region used in comparisons between raw input and RCAN predictions.

#### Expansion Microscopy (ExM)

Expansion microscopy was performed as described^26^. Immunolabeled U2OS cells were post-fixed in 0.25% Glutaraldehyde/1X PBS for 10 minutes at RT and rinsed three times in 1X PBS. Fixed cells were incubated with 200 μL of monomer solution (19% (wt/vol) Sodium acrylate (Sigma, 408220), 10% (wt/vol) Acrylamide (Sigma, A3553), 0.1% (wt/vol) N,N-Methylenebis(acrylamide) (Sigma, 146072) in 1X PBS) for 1 minute at RT. To start gelation, the monomer solution was replaced by fresh monomer solution containing 0.2% (vol/vol) Ammonium Persulfate (Thermofisher, 17874) and 0.2% (vol/vol) Tetramethylethylenediamine (Thermofisher, 17919). Gelation was allowed to proceed 40 minutes at RT, and the resulting gel was digested in 1 mL of digestion buffer (0.8M guanidine hydrochloride and 0.5% Triton X-100 in 1X TAE buffer) by Proteinase K (0.2mg/mL, Thermofisher, AM2548) for 1 hour at 45°C. After digestion, gels were expanded in 5mL of pure water (Millipore, Direct-Q 5UV, ZRQSVR5WW), and fresh water exchanged 3-4 times every 15 minutes.

#### Pre-ExM and post-ExM on the same cell

To compare images between pre- and post-ExM, the same group of cells needs to be located and imaged before and after ExM (**Supplementary Fig. 16**). After initial antibody (**Supplementary Table 1**) and DAPI staining, the pre-ExM cells were imaged under a wide field microscope with a 20X air objective (Olympus, UPlanFL N, 0.5 NA). Based on the DAPI signal, the nuclear shape, diameter, and distribution pattern of selected cells can be recorded, a useful aid in finding the same cells again if post-ExM images are acquired on the wide field microscope. The coarse location of a group of cells was marked by drawing a square with a Sharpie marker underneath the coverslip. The marked cells are then imaged on our home-built instant structured illumination microscope (iSIM^3^) before and after ExM in later steps. Before expansion, the marked region was imaged on iSIM with a 60X, NA 1.2 water objective (Olympus, PSF grade) to acquire pre-ExM data. The correction collar was adjusted to the 0.19 setting, which was empirically found to minimize spherical aberration. After ExM, a square portion of expanded gel was cut out, based on the marked region drawn underneath the cover glass, then remounted on a poly-L-lysine coated glass bottom dish (MatTek, P35G-1.5-14-C) and secured by depositing 0.1% low melt agarose around the periphery of the gel. To create the coated glass bottom dish, we applied poly-L-lysine (0.1% in water, Sigma, P892) for 30 minutes at room temperature, rinsed three times with pure water, and air dried. The same group of cells was then found on the wide field microscope using the DAPI stain and the 20X air objective. By comparing to the wide field DAPI image acquired before expansion, coarse estimation of the expansion factor as well as potential cell distortion/damage can be assayed. Finally, another square was drawn underneath the coverslip to locate the expanded cells, which were then imaged on the iSIM with the same objective and correction collar settings for post-ExM image acquisition.

#### Attempting to register pre- and post-expansion data

Pre- and post-expansion images were registered using the landmark registration module in 3D Slicer^36^ (http://www.slicer.org/). Landmark-based registration in 3D Slicer is an interactive registration method that allows the user to view registration results and manipulate landmarks in real time. We first rescaled the pre-expansion images according to the estimated expansion factor in the X, Y, and Z axes. During the registration process, pre-expansion images were used as fixed volumes and post-expansion images were used as moving volumes. Pre- and post-expansion images were coarsely aligned by affine registration based on 2-3 manually selected landmarks. Image registration was further refined using thin plate spline registration by interactively manipulating the landmarks. Finally, a transformation grid was generated to transform the post-expansion images to the pre-expansion images (**Supplementary Fig. 17**).

#### Estimating expansion factor

Pre- and post-expansion mitochondrial and microtubule data were inspected in 3D Slicer and registered with landmark-based registration as described above. Apparent distances between feature points were manually measured and ratioed to obtain the local expansion factor, which varied between 3.1-3.4 for mitochondria and 3.9-4.1 for microtubules (**Supplementary Fig. 15**). Based on this analysis we used a value 3.2 for mitochondria and 4.0 for microtubules in all downstream processing.

#### Stage scanning with iSIM

To rapidly tile multiple iSIM image fields to capture large expanded samples, we added a stage scan function into our control software, available on request from the authors. In the software, a step size of 0 to 150 μm can be selected for both horizontal (X) and vertical (Y) directions. We set this step size to be ∼70 μm, a value smaller than the field of view to ensure that each image had at least 20% overlap with adjacent images for stitching. We used up to 100 steps in both directions. The stage scan experiment was performed in a “zigzag” format (adjacent rows were scanned in opposite directions) to avoid large movements and maintain sample stability. At each stage position, 3D stacks were acquired. Stacks were stitched in Imaris Stitcher (Bitplane).

#### Generating synthetic pre-expansion data

To first order, we can interpret the post-expansion image as enlarging the object *s* by an expansion factor *M* and blurring with the system PSF, *sPSF*:

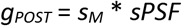

where *s_M_* is the expanded object, *g_POST_* is the post-expansion image of the expanded object, and * is the convolution operation. Similarly, if we upsample the pre-expansion image by a factor *M* we can approximate it as

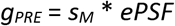

where *ePSF* is *sPSF* enlarged *M* times. We seek to express *g_PRE_* in terms of *g_POST_*, thus obtaining an estimate of *g_PRE_* in terms of the measured post-expansion image.

Fourier transforming (FT) both equations, dividing to eliminate the object spectrum, and rearranging terms, we obtain

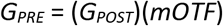

where *mOTF* is a modified OTF equivalent to the ratio of the OTFs corresponding to *ePSF* and *sPSF*, i.e. *mOTF* = FT(*ePSF*) / FT(*sPSF*). To avoid zero or near zero division in this calculation, we set the amplitude of FT(sPSF) to 1 beyond the cut-off frequency of *sPSF*. Finally, inverse Fourier transforming yields a synthetic estimate of *g_PRE_*.

We improved this estimate by also modifying the background and noise levels to better match experimental pre-expansion images, computing the SSIM between the synthetic image and the experimental pre-expansion image as a measure of similarity. We tried to maximize the SSIM by (1) laterally and axially modifying the modelled *sPSF* so that the FWHM value is equal to the FWHM measured with 100 nm beads and resolution-limited structures in the experimental images; (2) modifying the background level, i.e., adding or subtracting a constant value; and (3) adding Gaussian and Poisson noise. We optimized these parameters in a range +/− 15 % of the values derived from experimental pre-expansion data (2-3 pre-expansion images that could be reasonably well registered to corresponding post-expansion data), and then applied these optimized parameters for all synthetic data. Finally, we performed a visual check before deconvolving the synthetic data and post-expansion data in preparation for RCAN training. 15 iterations of Richardson-Lucy deconvolution were applied, using *sPSF* for the expanded images and the modified *ePSF* for the synthetic data. These steps are shown in **Supplementary Fig. 19**.

#### Estimating signal-to-noise (SNR) ratio in experiments

We assumed a simple model for per-pixel SNR, accounting for Poisson noise arising from the signal and read noise from the camera. After subtracting a constant background offset (100 counts) and converting the digital signal in each pixel to photons using the manufacturer-supplied conversion factor (0.46 photoelectrons/digital count), we used

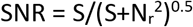

where S is the observed, background-corrected signal in photoelectrons, and N_r_ the read noise (1.3 electrons from the manufacturer).

#### Spherical simulations

For the images in **Fig. 2** and images and analysis in **Supplementary Fig. 5, 8, 9** the simulated ground-truth images consisted of spheres seeded at random locations and with random size and intensity, generated with ImgLib2^37^. The maximum radius of the spheres was set at 3 pixels and the intensity range set to 1000 to 20000. We generated a set of 30 such images with size of 256×256×256. Ground truth (GT) images were generated by blurring this set of 30 images with the iSIM PSF (simulated as the product of the excitation and emission PSFs, generated in PSF generator (http://bigwww.epfl.ch/algorithms/psfgenerator/) with an NA of 1.42 and wavelengths 488 nm and 561 nm, respectively). Noisy phantom images were obtained by adding Gaussian noise (simulating the background noise of the camera in the absence of fluorescence) and Poisson noise (proportional to the square root of the signal) to the GT images. The 2x, 3x and 4x blurred noiseless phantom images were obtained by blurring the initial 30 images with a kernel 2x, 3x and 4x the size of the iSIM PSF.

#### Estimating spatial resolution

The resolution measures in **Fig. 1d** were estimated by computing the FWHM as a measure of apparent size of a subdiffractive object (microtubule width). However, all other resolution estimates were based on decorrelation analysis^24^. This method estimates average image resolution from the local maxima of a series of decorrelation functions, providing an estimated resolution that corresponds to the highest spatial frequency with sufficient SNR, rather the Abbe resolution limit.

There are four main steps in the algorithm. First, the Fourier transform of the input image *I*(*k*) and its normalized version *I_n_*(*k*) are cross-correlated using Pearson correlation, producing a single value between 0 and 1, denoted *d*. Second, the normalized Fourier transform *I_n_*(*k*) is repeatedly filtered by a binary circular mask with different radius *r* ∈ [0,1] (here *r is* expressed as a normalized spatial frequency), and the cross-correlation between *I*(*k*) and each filtered *I_n_*(*k*) is recalculated, yielding a decorrelation function *d*(*r*). This decorrelation function exhibits a local maximum of amplitude *A_0_* that indicates the spatial frequency *r_0_* of best noise rejection and signal preservation ratio. Third, the input image is repeatedly filtered with different Gaussian high-pass filters to attenuate the energy of low frequencies. For each filtered image, another decorrelation function is computed, generating a set of [*r_i_, A_i_*] pairs, where *r_i_* and *A_i_* are the position and amplitude of the local maximum, respectively. Last, the most suitable peak position (i.e., selected from the *r_i_*) is selected as the estimate of resolution. In the original algorithm^24^, two choices are used and validated in many applications - (1) the peak corresponding to the highest frequency (i.e., the maximum *r_i_* value); (2) the peak corresponding to the highest geometric mean of *r_i_* and *A_i_*.

However, we found that both criteria often failed when using them on our images, i.e., the estimated resolution was often a value much beyond the theoretic resolution limit. Plotting [*r_i_, A_i_*] pairs shows three phases: *A_i_* first increases in phase I, then gradually decreases in phase II, and finally increases again in phase III (**Supplementary Fig. 11a**). Resolution values in phase III exist due to digital upsampling of the pixel size, but are not reliable, as they extend past the Abbe limit. We thus modified the algorithm by (1) setting a theoretical resolution limit in computing the decorrelation functions; and (2) adopting a new criterion to determine the resolution estimate. Our new criterion finds the local minimum of *A_i_* to locate *r_i_* at the transition between phase II and phase III, which provides a reliable resolution estimate that is robust to changes in pixel size. We validated this strategy on a microtubule image with 1x, 1.5x and 3x digital upsampling (**Supplementary Fig. 11b**), finding that our criterion gave identical estimates of spatial resolution in each case.

For estimating the lateral and axial resolution in our data (input, ground truth, and deep learning outputs), we first interpolated the stacks along the axial dimension to achieve isotropic pixel size. Then we performed our modified decorrelation analysis on a series of xy slices to obtain lateral resolution estimate (with mean and standard deviations derived from the slices). For axial resolution, we implemented sectorial resolution estimate^24^ on a series of xz slices, where the binary circular mask was replaced with a sectorial mask (22.5 degree opening angle, **Supplementary Fig. 11c**) that captured spatial frequencies predominantly along the z dimension.

### Confocal and STED microscopy

#### Sample preparation

Mouse embryonic fibroblasts (MEF) were grown in #1.5 glass-bottom dishes (MatTek, P35G-1.5-20-C) using DMEM (Gibco, 10564011) supplemented with 10% FBS (Quality Biological, 110-001-101HI).

For microtubule and nuclear pore samples, we fixed and permeabilized cells with −20°C methanol (Sigma-Aldrich, 322415) for 10 min at −20°C. Samples were rinsed and blocked for 1 hour with 1x Blocker BSA (ThermoFisher Scientific, 37525) and incubated overnight with 1:500 dilution of primary rabbit anti-alpha tubulin (Abcam, ab18251) and mouse anti-nuclear pore complex (Abcam, ab24609) antibodies in 1x Blocker BSA at 4°C. Samples were washed three times for 5 minutes with 1x Blocker BSA. After the last washing step, we fluorescently labeled samples by incubation with 1:500 dilution of secondary Alexa Fluor 594 goat anti-mouse (ThermoFisher Scientific, A-11005) and ATTO 647N goat anti-rabbit (Sigma-Aldrich, 40839) antibodies in 500 μL of 1x Blocker BSA for 4 hours at room temperature. Samples were washed four times for 5 minutes with 1x Blocker BSA. After final washing, samples were mounted in glass-bottom dishes using 90% Glycerol (Sigma-Aldrich, G2025) in PBS (KD Medical, RGF-3210).

For SiR-DNA imaging, we used live MEF cells, grown as before, and MEF cells fixed with 4°C 4% formaldehyde (Sigma-Aldrich, 252549) in PBS for 20 minutes at room temperature. Sample labelling was performed with the SiR-DNA kit (Spirochrome, SC007) following the manufacturers protocol: https://spirochrome.com/documents/202003/datasheet_SPY650-DNA_202003.pdf. Fixed samples were mounted as before.

#### Imaging

We acquired 33 matched sets of confocal/STED volumes for microtubule- and nuclear pore complex-labeled samples. For these experiments all images were acquired using a Leica SP8 3X STED microscope, a white-light laser for fluorescence excitation (470-670nm), a Leica HyD SMD time-gated PMT, and a Leica 100x (1.4 N.A.) STED White objective (Leica Microsystems, Inc.). ATTO 647 was excited at 647 nm and emission was collected over a bandwidth of 657-700 nm. Alexa Fluor 594 was imaged with 580 nm excitation, and emission was collected over a bandwidth of 590-650 nm. All images (both confocal and STED) were acquired with a pinhole size of 0.7 A.U., a scan speed of 600 Hz, a pixel format of 1024 x 1024 (pixel sizes of 25 nm), a 6-slice z-stack acquired at an interslice distance of 0.16 μm, and time gating on the HyD SMD set to a time range of 0.7-6.5 ns. The STED images for both labels were acquired with depletion at 775nm laser (pulsed at 80 MHz) at a power of 105 mW at the back aperture for ATTO 647-labeled microtubules (25% of full power) and 85 mW at the back aperture for Alexa Fluor 594 labeled nuclear pore complexes (20% of full laser power). Fluorescence excitation for STED imaging was set to 4x and 1.5x the confocal excitation power levels for ATTO 647 and Alexa Fluor 594 respectively. For ATTO 647, HyD SMD gain was set to 100% for confocal and STED imaging. For Alexa Fluor 594, HyD SMD gain was set to 64% for confocal imaging and 100% for STED imaging. For both colors, confocal images were acquired with a 2-frame line average and STED images were acquired with a 2-frame line average combined with 2-frame integration.

SiR-DNA labelled MEF cells were imaged both in the fixed (confocal and STED) and live-cell (confocal only) mode. Low SNR confocal and high-quality STED image replicates were taken on similar fixed samples (35 data sets) to train a deep-learning model to apply to the live cell confocal data. Low-excitation level (thus low SNR) live cell confocal images were followed over time to capture cell division. For these experiments, the same microscope hardware listed above was used but scanned in the resonant mode (to afford more rapid imaging capable of capturing cell division). For live cell confocal stacks of 25 or more slices (interslice distance of 0.16 μm) were taken approximately every minute (2s/frame) continuously for a period of ∼30-45 minutes. Images were taken with a scan rate of 8000 Hz, 8 line-average, a pinhole set to 1 A.U, 647nm excitation (5% of total laser power), an emission bandwidth of 657-637nm, and at a pixel size of 25 nm at a format of 2048 x 2048. For the fixed cell experiments the confocal settings were the same except that line averaging was set to 16, the frame rate was 6s/frame, the excitation power at 647nm was set to 0.1% total laser power (to approximately match the SNR in the live cell data), and only one z-stack was taken. STED experiments were the same, except that 647 nm excitation was set to 1.5% and the depletion power at 775 nm was 7.5% (approximately 35 mW at the back aperture). Time gating windows on the HyD SMD was set to 0.3 to 6.5 ns or 0.7 to 6.5ns for the confocal and STED experiments, respectively. For live experiments, temperature was set to 37°C using a culture dish heater and temperature control unit (DH-35 and TC-344B, Warner Instruments, Hamden, CT) and an objective heater (Bioptechs, Butler, PA).

#### Deconvolution

Huygens Professional (version 19.1, Scientific Volume Imaging, Hilversum, The Netherlands) was used to deconvolve some confocal images. All deconvolution was based on idealized point spread functions, using the classic maximum likelihood estimation (CMLE) deconvolution algorithm. In some cases, the object stabilizer module was used to compensate for drift and minor mechanical instabilities.

#### Data availability

The data that support the findings of this study are available from the corresponding author upon reasonable request.

#### Code availability

The code used in this study will be available as Supplementary Software. We plan to upload software and supplementary data to GitHub in a finalized version of this manuscript.

