## Supplementary Information for "Three-dimensional residual channel attention networks denoise and sharpen fluorescence microscopy image volumes"

**Supplementary Video Captions**

**Supplementary Video 1, iSIM imaging at high SNR rapidly bleaches the sample.** Live U2OS cells expressing pShooter pEF-Myc-mito-GFP were imaged with high intensity illumination every 5.6 s, leading to obvious photobleaching. Maximum intensity projections are shown. Scale bar: 5 μm. See also **Fig. 1e**.

**Supplementary Video 2, RCAN enables super-resolution imaging over thousands of volumes.** Live U2OS cells expressing pShooter pEF-Myc-mito-GFP were imaged with low intensity illumination every 5.6 s, recording 2600 volumes. Left: raw iSIM input; Right: RCAN prediction. Maximum intensity projections are shown. Scale bar: 5 μm. See also **Fig. 1e**.

**Supplementary Video 3, Two-color volumetric image restoration with RCAN.** Live U2OS cells expressing mApple-Lamp1 (labeling lysosomes, red) and pShooter pEF-Myc-mito-GFP (labeling mitochondria, green) were imaged every 5.1 s for 300 volumes. Left: raw iSIM input; Right: RCAN prediction. Maximum intensity projections are shown. Scale bar: 5 μm. See also **Fig. 1f**.

**Supplementary Video 4, Axial views of 2x blurred input, and network predictions.** Noiseless spherical phantoms are blurred with a kernel two-fold larger than the PSF (upper left) and used as an input to CARE, SRResNet, ESRGAN, and RCAN as shown. Axial reslices through stacks are shown. See also **Fig. 2**.

**Supplementary Video 5, Axial views of 3x blurred input, and network predictions.** Noiseless spherical phantoms are blurred with a kernel three-fold larger than the PSF (upper left) and used as an input to CARE, SRResNet, ESRGAN, and RCAN as shown. Axial reslices through stacks are shown. See also **Fig. 2**.

**Supplementary Video 6, Axial views of 4x blurred input, and network predictions.** Noiseless spherical phantoms are blurred with a kernel four-fold larger than the PSF (upper left) and used as an input to CARE, SRResNet, ESRGAN, and RCAN as shown. Axial reslices through stacks are shown. See also **Supplemnetary Fig. 9**.

**Supplementary Video 7, RCAN denoises and improves resonant confocal recordings of dividing cells**. Live MEF cells were stained with SiR-DNA and volumetrically imaged with resonant confocal microscopy every 58 s. Left: raw resonant confocal data; Right: RCAN prediction. Maximum intensity projections are shown. Scale bar: 5 μm. See also **Fig. 3e, f**.

**Supplementary Video 8, RCAN denoises and improves resonant confocal recordings on additional nuclei**. Live MEF cells were stained with SiR-DNA and volumetrically imaged with resonant confocal microscopy every 58 s. Left: raw resonant confocal data; Right: RCAN prediction. Top/Bottom: XY/XZ maximum intensity projections. Scale bar: 5 μm.

**Supplementary Video 9, Synthetic input, RCAN prediction, and ground truth for mitochondrial images based on expansion microscopy training.** Mitochondria in fixed U2OS cells were immunolabeled against Tomm-20, expanded, synthetically degraded, and used to train an RCAN model. Z stacks are shown. Left: synthetic input ('RAW'), meant to mimic noisy, pre-expanded iSIM deconvolution; middle: RCAN prediction; right: deconvolved, expanded ground truth data. Scale bar: 5 μm. See also **Fig. 4b**.

**Supplementary Video 10, Live-cell mitochondrial RCAN prediction based on expansion microscopy training.** Live U2OS cells expressing mEmerald-Tomm20 were volumetrically imaged every 10 s with iSIM and deconvolved images input into a trained RCAN model. Top/bottom: lateral/axial maximum intensity projections. See also **Fig. 4d**.

**Supplementary Video 11, High magnification axial view, comparing deconvolved iSIM input and RCAN prediction.** Live U2OS cells expressing mEmerald-Tomm20 were volumetrically imaged every 10 s with iSIM and input into the trained RCAN model. Here an axial view corresponding to that in **Fig. 4d** is shown, comparing deconvolved iSIM input and RCAN prediction. Scale bar: 2 μm.

**Supplementary Video 12, Live-cell microtubule RCAN prediction based on expansion microscopy, axial views.** Live Jurkat T cells expressing EMTB-3XGFP were volumetrically imaged every 12.3 s with iSIM and input into a trained RCAN model. Axial maximum intensity projections are shown for deconvolved iSIM (top) and RCAN (bottom). Scale bar: 2 μm. See also **Fig. 4e**.

**Supplementary Video 13, Live-cell microtubule RCAN prediction based on expansion microscopy, lateral views.** Live Jurkat T cells expressing EMTB-3XGFP were volumetrically imaged every 12.3 s with iSIM and input into a trained RCAN model. Lateral maximum intensity projections corresponding to the lateral region of interest in **Fig. 4e** are shown. The 'FIRE' color map in ImageJ is used to improve visibility of dim filaments near the cell boundary. Scale bar: 1 μm. See also **Fig. 4e**.

**Supplementary Video 14, Additional example of microtubule dynamics in another Jurkat T cell.** Live Jurkat T cells expressing EMTB-3XGFP were volumetrically imaged every 12.3 s with iSIM and input into a trained RCAN model. Rotational maximum intensity projections are shown for deconvolved iSIM (left) and RCAN (right).

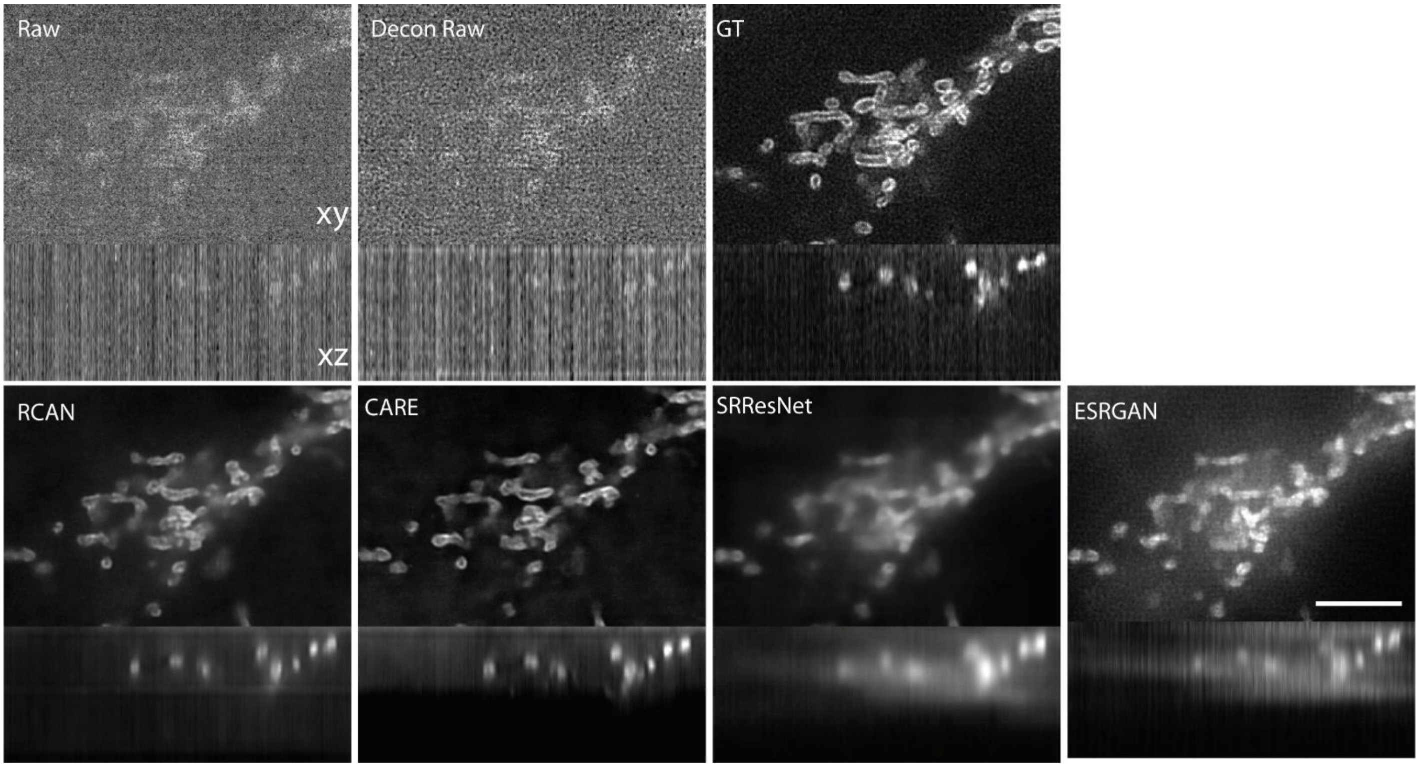


**Supplementary Fig. 1, Denoised neural network output outperforms direct deconvolution of noisy input.** Fixed U2OS cells expressing mEmerald-Tomm20 were imaged on the iSIM at low (Raw) and high (ground truth, GT) power. We also deconvolved the raw input (Decon Raw) and computed the RCAN, CARE, SRResNET and ESRGAN predicions. Lateral (xy) and axial (xz) slices are shown. Scale bar: 5 μm. See also **Fig. 1b**.

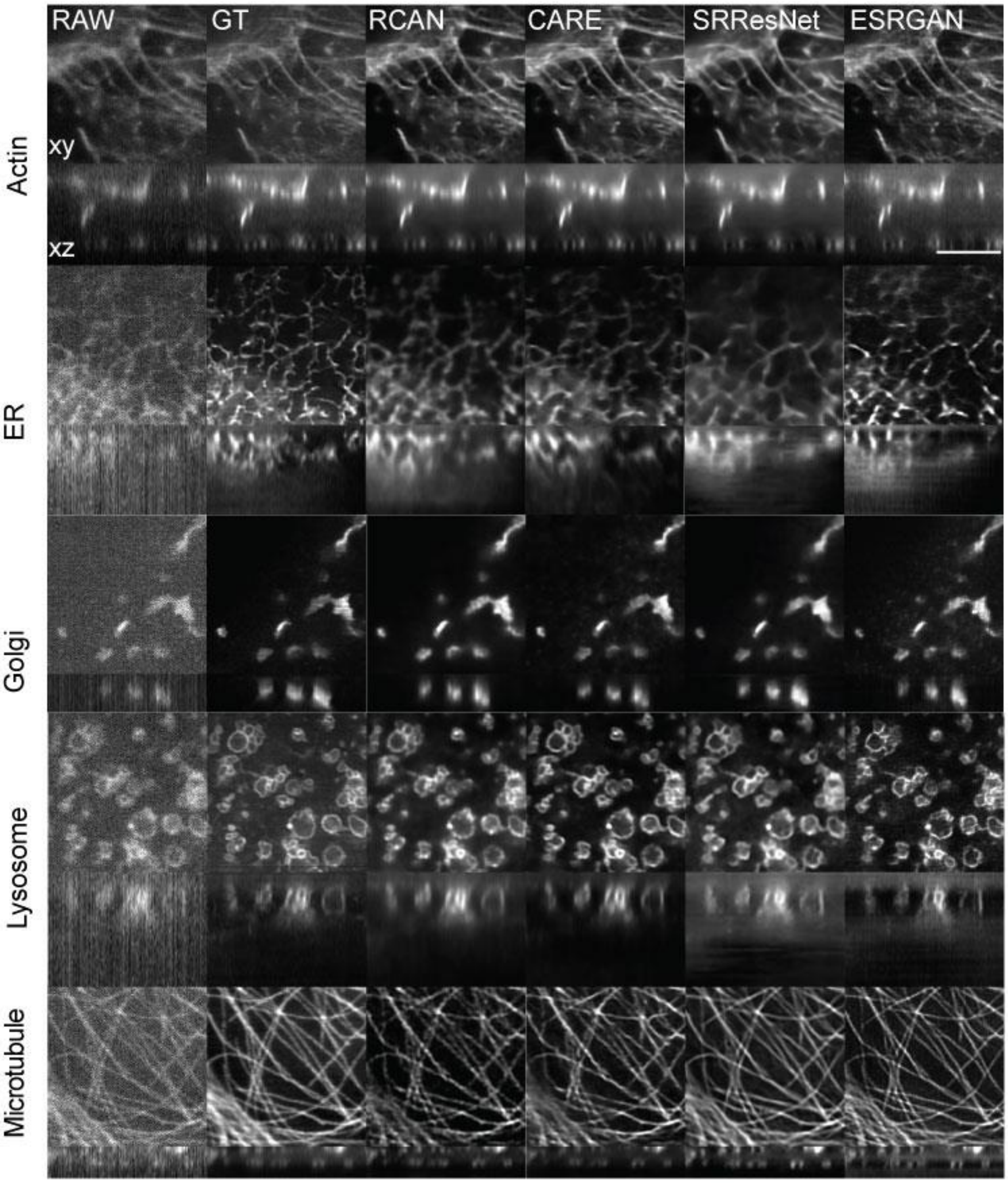


**Supplementary Fig. 2**, **Organelle denoising by different networks.** Rows indicate organellese, stained with Alexa Fluor 488 Phalloidin (actin); ERmoxGFP (endoplasmic reticulum, ER); GalT-GFP (golgi); LAMP1-mApple (lysosomes); and Mouse-α-Tubulin primary antibody, Donkey- α-Mouse Biotin secondary antibody and Alexa Fluor 488 Streptavidin (microtubules). Columns indicate low SNR raw input data, high SNR deconvolved ground truth, RCAN prediction, CARE prediction, SRResNet prediction, and ESRGAN prediction. Lateral (top) and axial (bottom) slices are shown for each row. Scale bar: 5 μm. See **Supplementary Table 4, Supplementary Fig. 3** for corresponding SSIM and PSNR values.

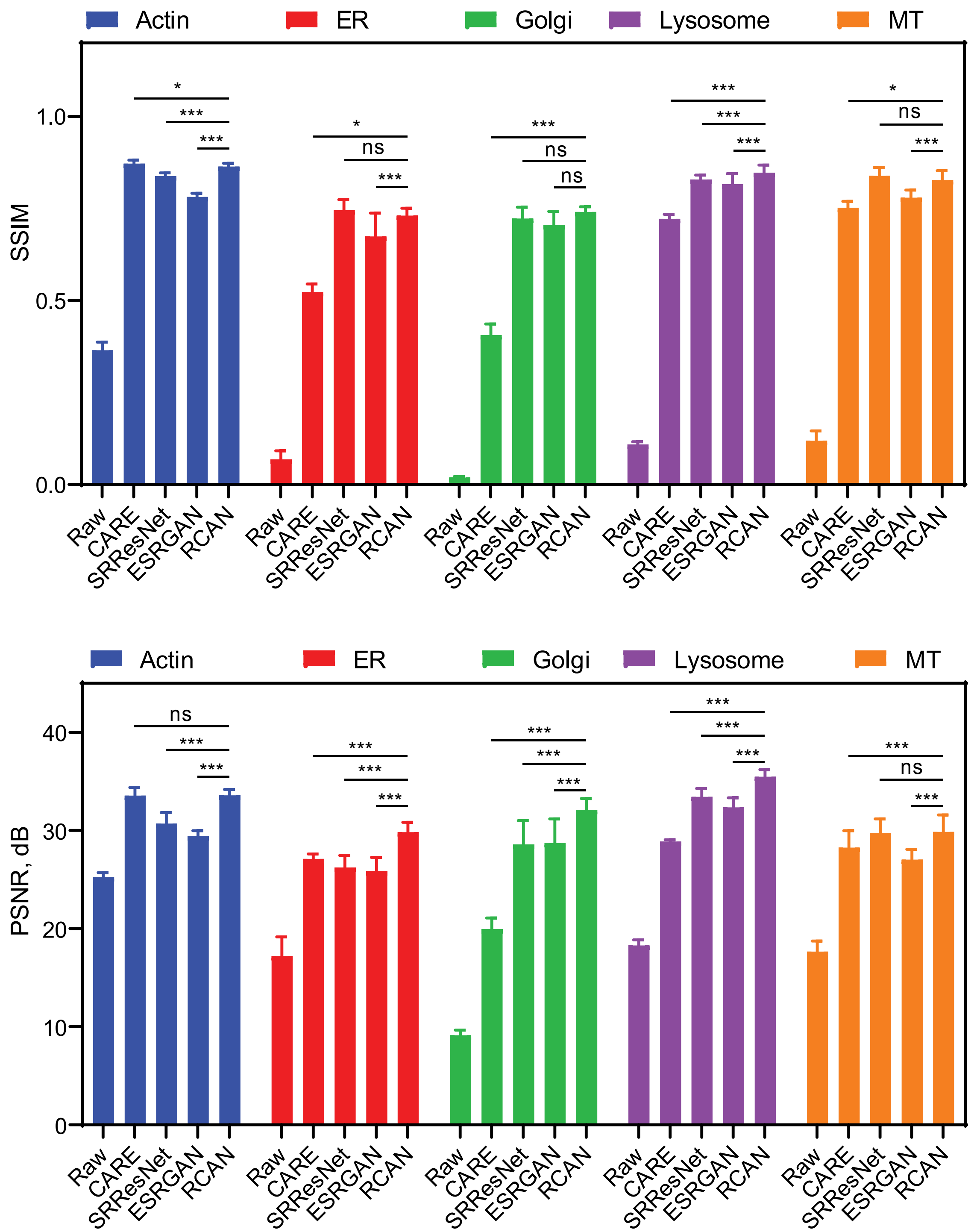


**Supplementary Fig. 3, Network denoising comparisons.** Structural similarity index (SSIM, top) and peak signal-to-noise ratio (PSNR, bottom) are shown, for raw data input and prediction from CARE, SRResNet, ESRGAN, and RCAN on fluorescent labels shown in **Supplementary Fig. 2**. Means and standard deviations from 10-21 imaging planes are shown (5 cells). ***: P<0.0001, *: P<0.05, ns: not significantly different. See also **Supplementary Table 3**.

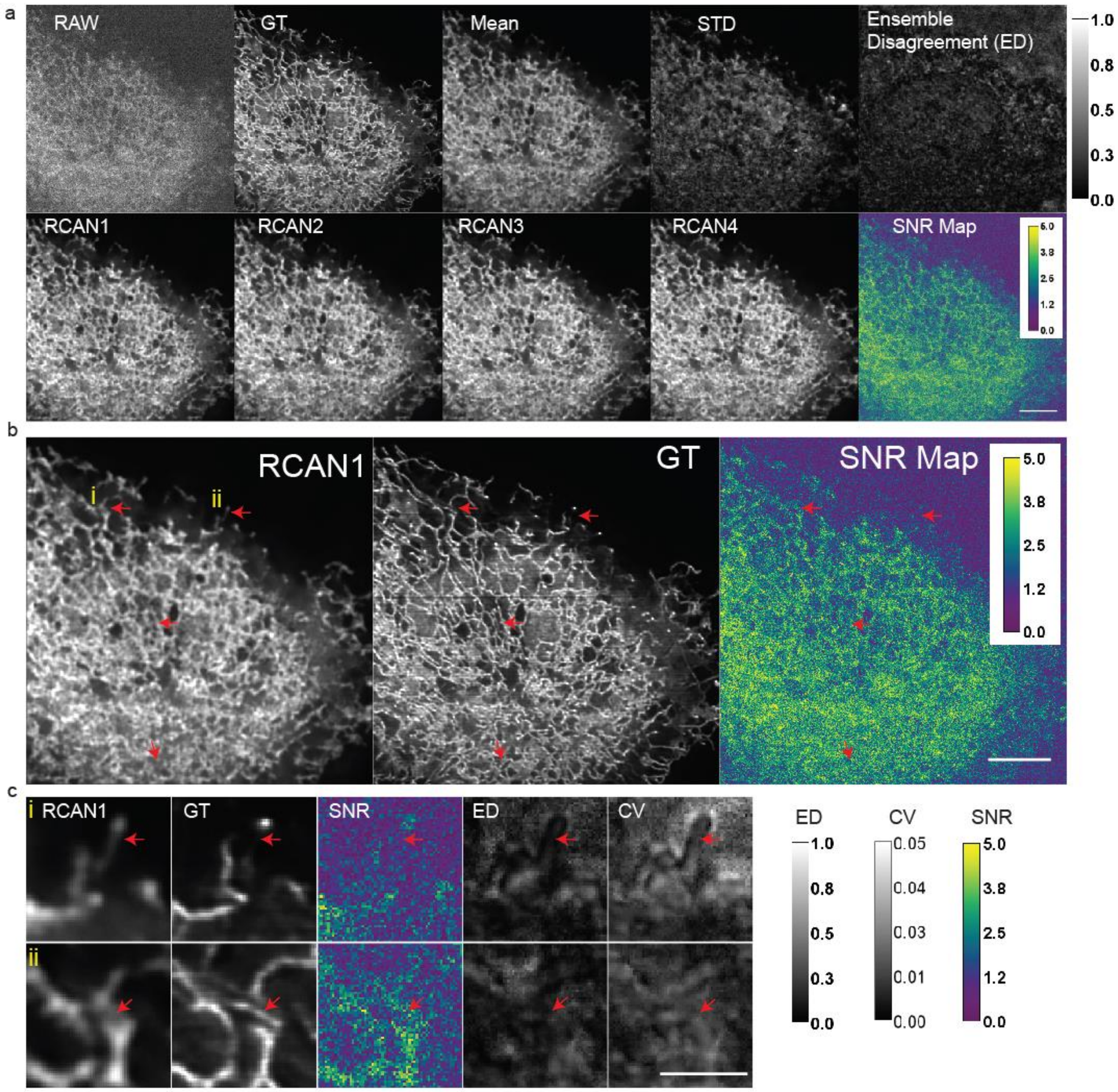


**Supplementary Fig. 4, RCAN performance metrics**. **a)** Top row: noisy input, high SNR ground truth, mean RCAN prediction, standard deviation RCAN prediction, and ensemble disagreement (ED). Bottom row: Four RCAN models, independently initialized and trained to denoise input, and SNR map based on noisy input image. Images show endoplasmic reticulum in fixed cells, labeled with ERmoxGFP and imaged with iSIM. **b)** Higher magnification view of first RCAN model, ground truth input, and input image SNR map. Red arrows highlight areas of discrepancy between RCAN output and ground truth. **c)** Higher magnification view of two areas (marked i, ii in **b**), illustrating network output, ground truth, input SNR, ED, and coeffient of variation (standard deviation of network output / mean of network output, from **a**). Scale bars: **a, b** 5 μm, **c** 2 μm.

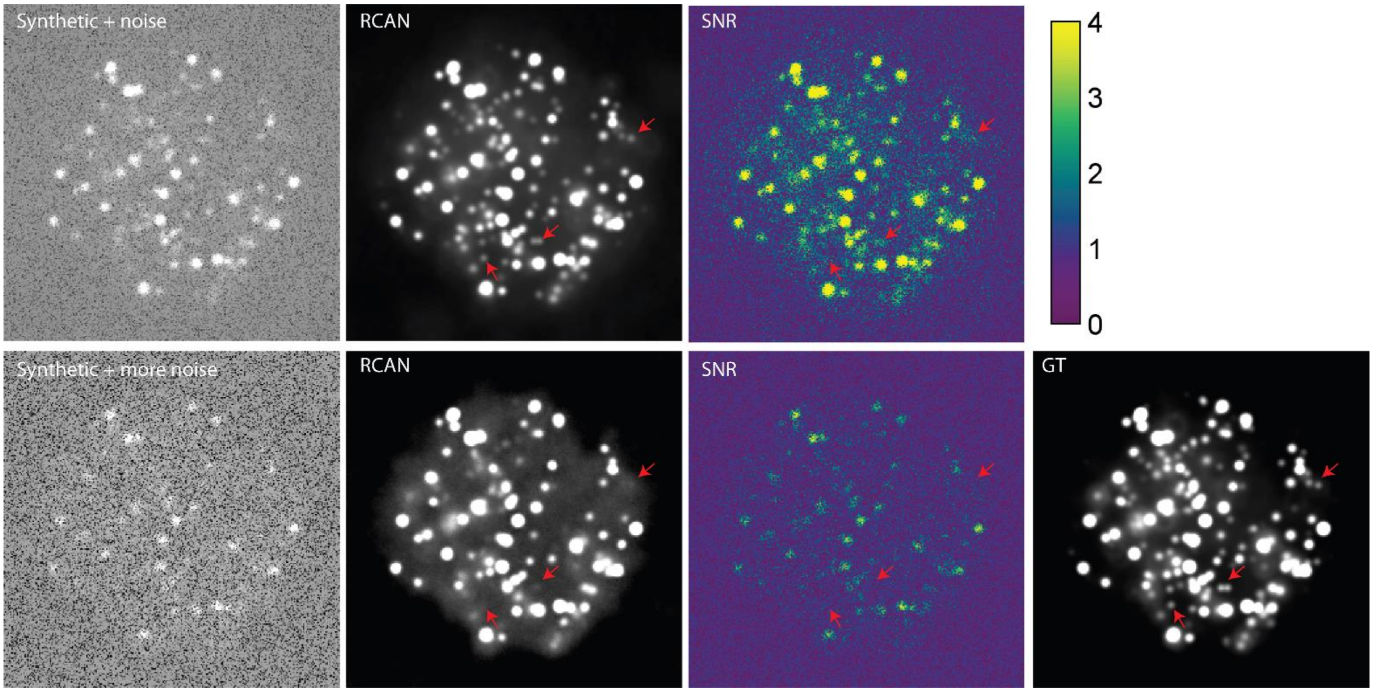


**Supplementary Fig. 5**, **Failure of RCAN prediction on noisy simulated data.** *Top*: Noisy synthetic sphere input, RCAN prediction, and per-pixel SNR signal-to-noise (SNR) map. *Bottom*: Same input as above but with more noise, RCAN prediction, SNR map, and ground truth (GT). Red arrows highlight features that are predicted using less noisy input but are not well predicted when using noisier input. See also **Methods**.

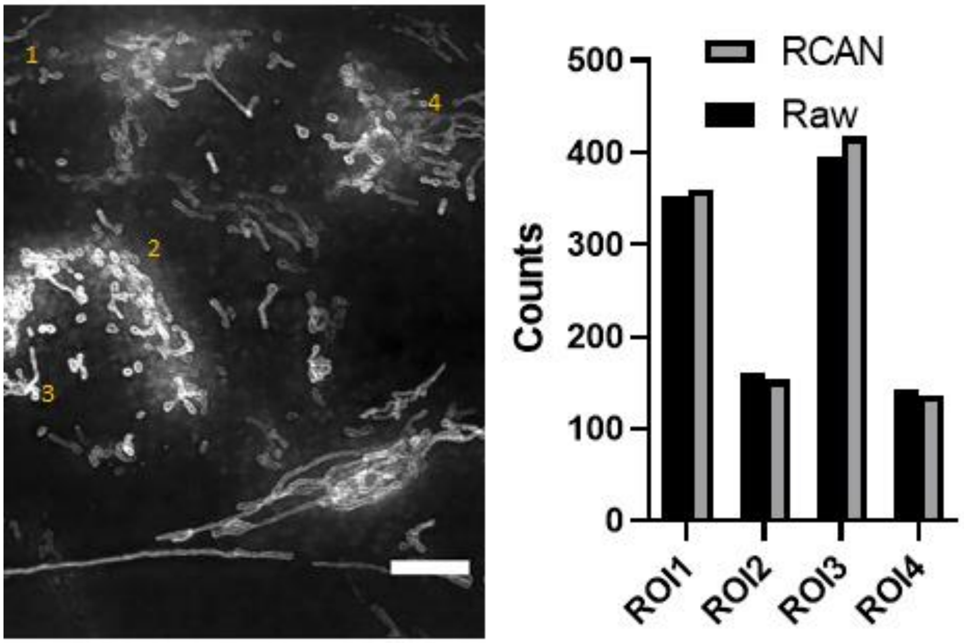


**Suppementary Fig. 6, RCAN preserves linearity**. *Left*: RCAN prediction (maximum intensity projection) for iSIM input image of fixed U2OS cells expressing mEmerald-Tomm20. Scale bar: 5 μm. *Right*: apparent intensity (counts in arbitrary units) at different regions of interest (average of 8x8 pixel regions, at locations indicated with orange numbers at left), in raw input data and RCAN prediction.

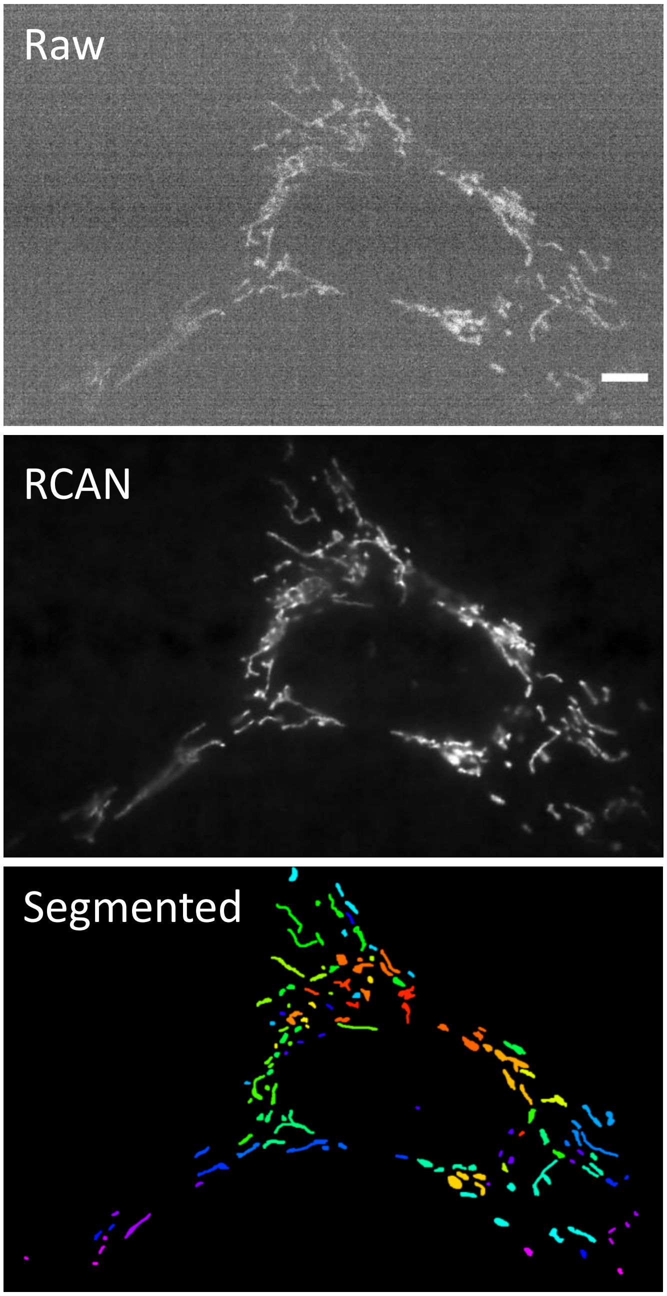


**Supplementary Fig. 7, Segmenting RCAN prediction.** Top: raw, noisy iSIM input. Middle: RCAN prediction. Bottom: manually segmented output, with segmented mitochondria shown in different colors. 153 disinct mitochondria were manually segmented in this volume, the first in the series shown in **Fig. 1e**. A single slice from the volume is shown here. Scale bar: 5 μm.

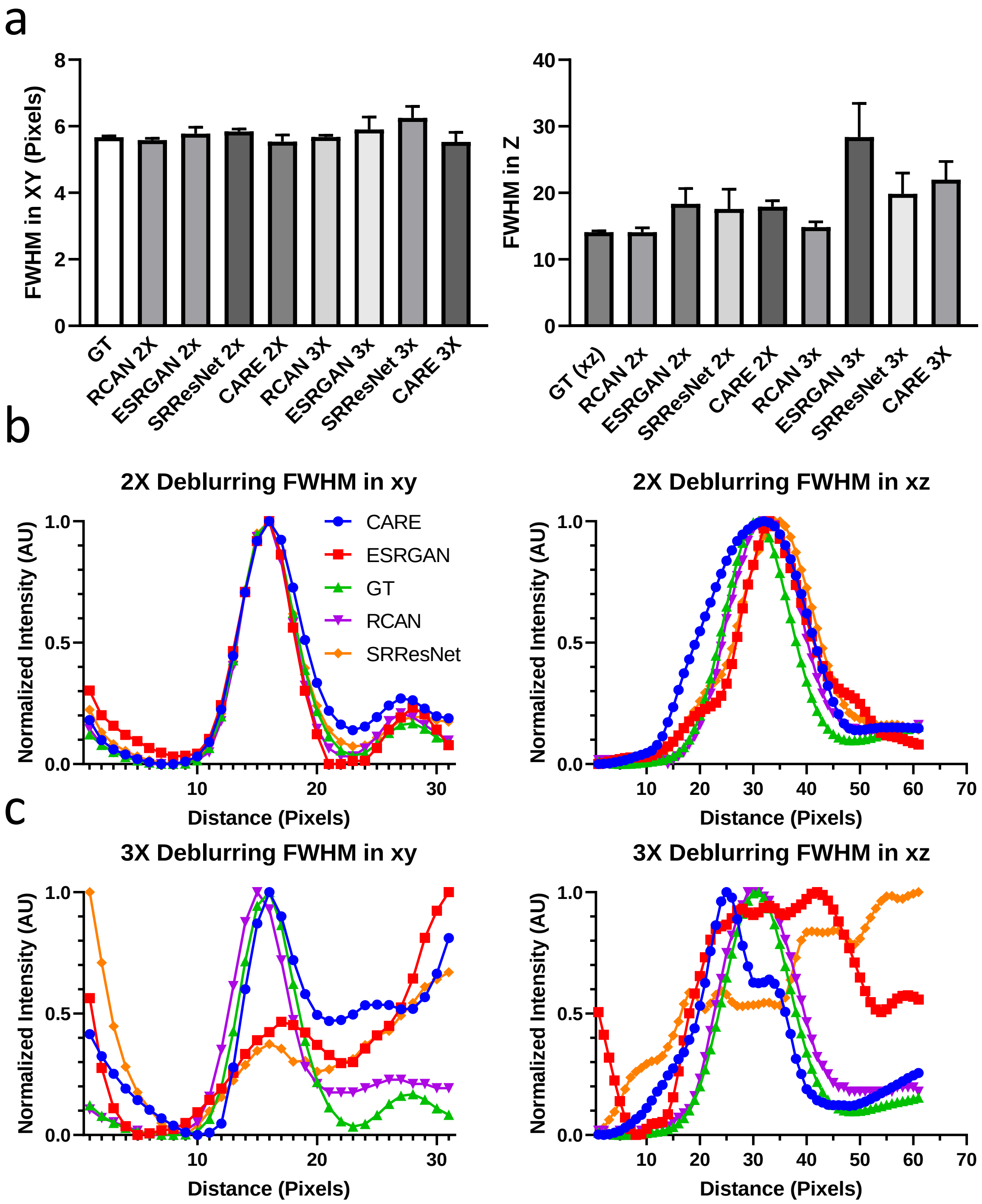


**Supplementary Fig. 8, Apparent size of subdiffractive beads after network restoration. a)** Lateral (left) and axial (right) full width at half maximum (FHWM) measurements for ground truth (GT), RCAN, ESRGAN, SRResNet, and CARE predictions. Means and standard deviations from measurements on 10-15 beads are shown. **b)** Lateral (left) and axial (right) line profiles corresponding to the bead shown in **Fig. 2b**, for data that has been blurred with a kernel two-fold larger than the PSF. **c)** As in **b)**, but for blurring with a kernel three-old larger than the PSF.

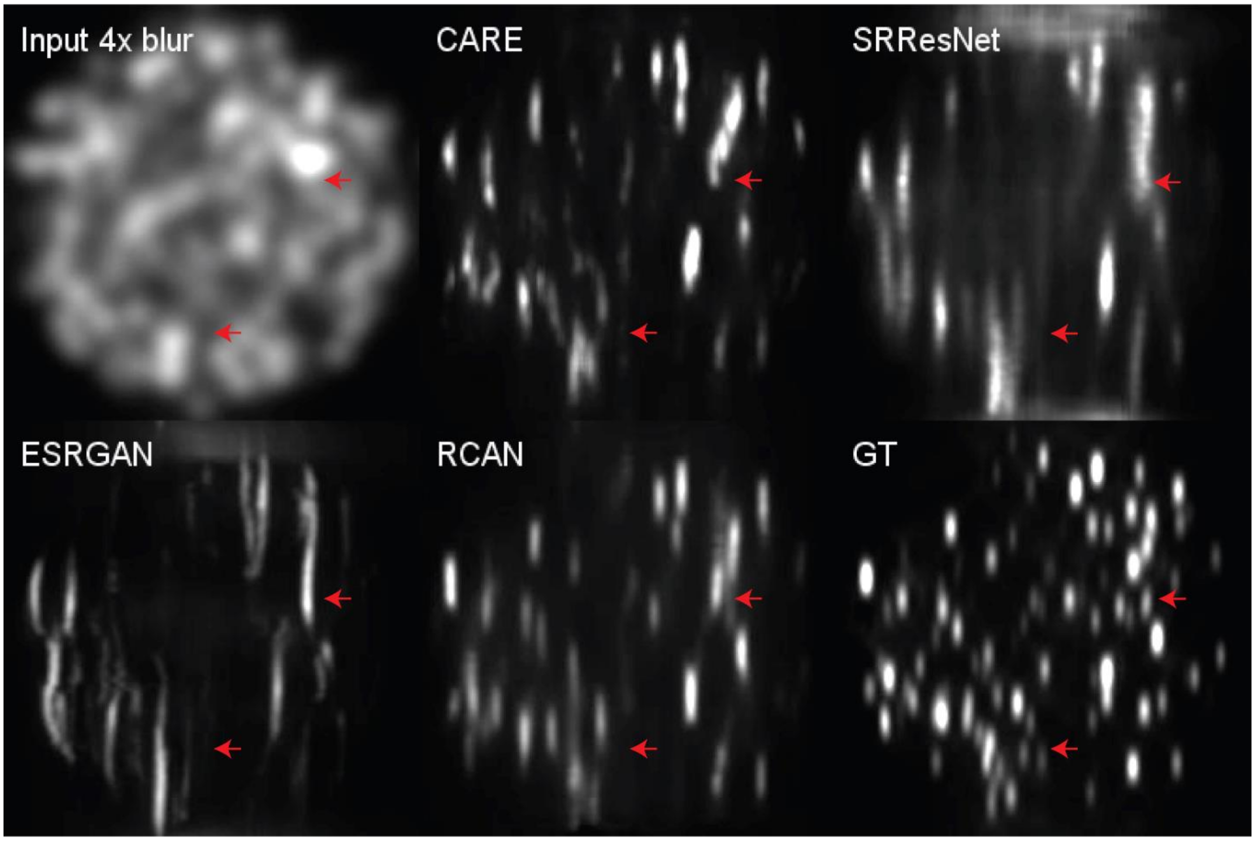


**Supplementary Fig. 9, Network performance at 4x blurring.** Data were blurred with a kernel four-fold larger than the PSF (Raw) and deblurred with CARE, SRResNet, ESRGAN, and RCAN. Axial reslices are shown, with red arrows highlighting areas of obvious discrepancy compared to ground truth (GT).

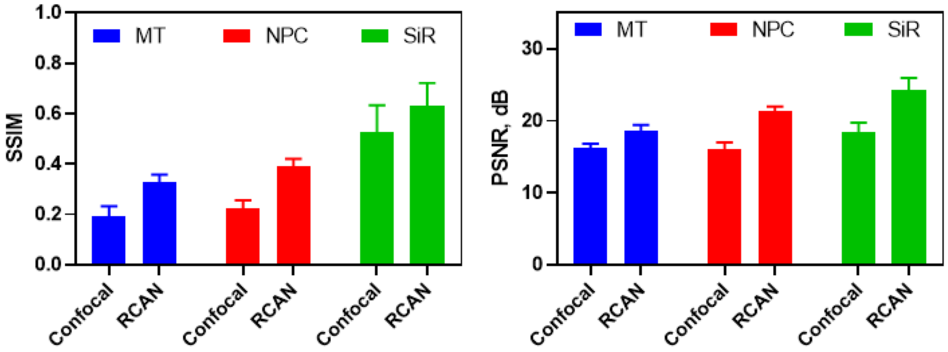


**Supplementary Fig. 10, SSIM and PSNR analyis for confocal to STED image restoration.** Structural similarity index (SSIM) and peak signal-to-noise ratio (PSNR) for datasets shown in **Fig. 3a-c.** Here 'Confocal' refers to the confocal input, and 'RCAN' is the prediction based on this input. $N$ = 20, 22, and 33 measurements were used for microtubule (MT), nuclear pore complex (NPC), and SiR-DNA (SiR), respectively. See also **Supplementary Table 4**.

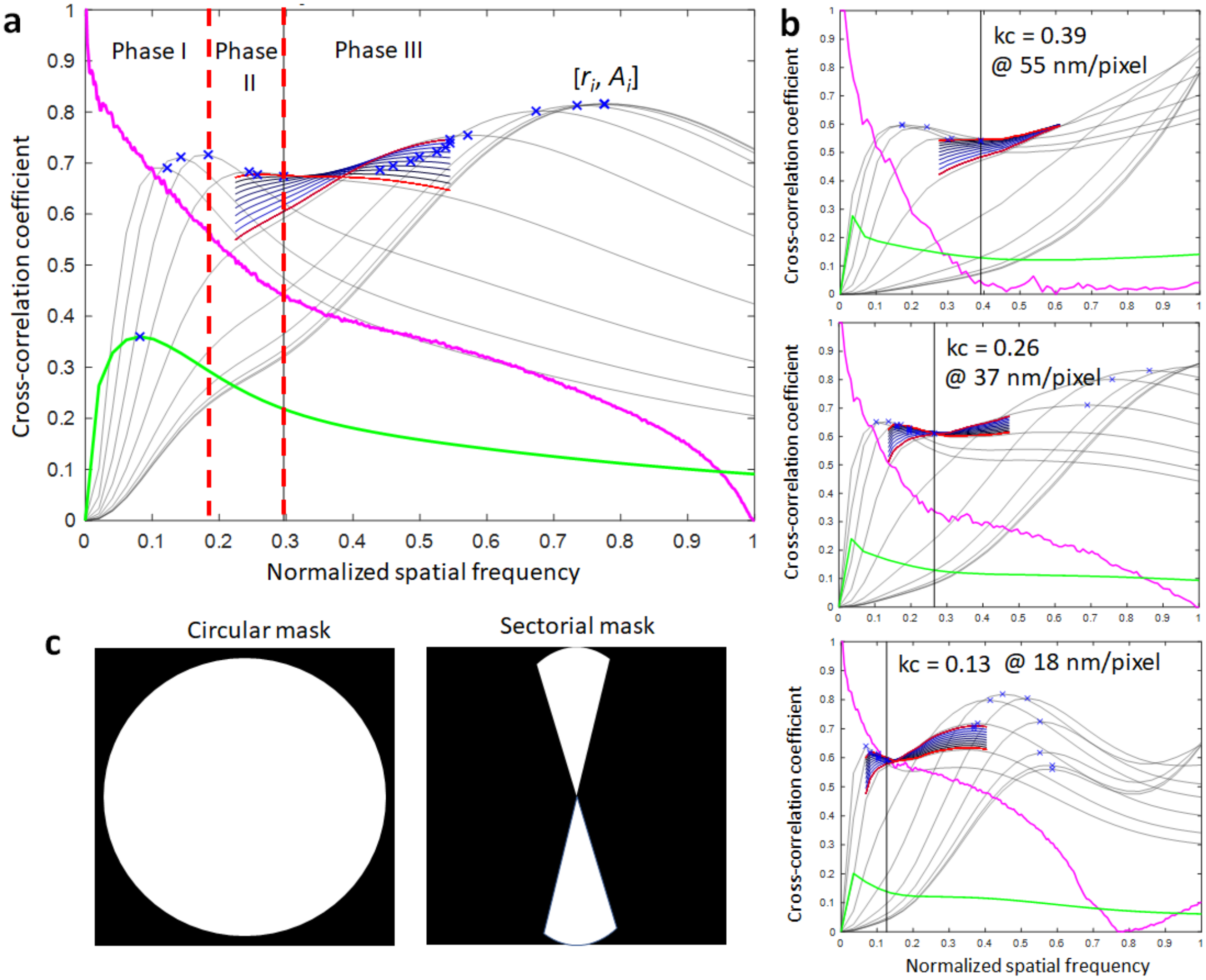


**Supplementary Fig. 11, Modified decorrelation analysis for resolution estimation. (a)** Example decorrelation functions computed for resolution estimation. Magenta line: radial average of log of absolute value of Fourier transform of the input image; green line: decorrelation function without any high-pass filtering; grey lines: decorrelation functions with high-pass filtering; blue cross markers: local maxima at $r_i$ with amplitude $A_i$. Red dashed lines split the pairs [$r_i$, $A_i$] into three phases: $A_i$ first increases in phase I, then gradually decreases in phase II, and finally increases again in phase III. **(b)** Plots of all decorrelation functions computed for a microtubule image without interpolation (55 nm/pixel, top panel), with 1.5x upsampling (37 nm/pixel, middle panel), and with 3x upsampling (18 nm/pixel, bottom panel). Our modified decorrelation method estimates the normalized cut-off frequency (kc) as 0.39, 0.26 and 0.13 for 1x, 1.5x and 3x upsampling, respectively. All values correspond to a spatial resolution of 282 nm and are thus robust to changes in pixel size. **(c)** Schematic of circular and sectorial mask for computing lateral and axial resolutions, respectively.

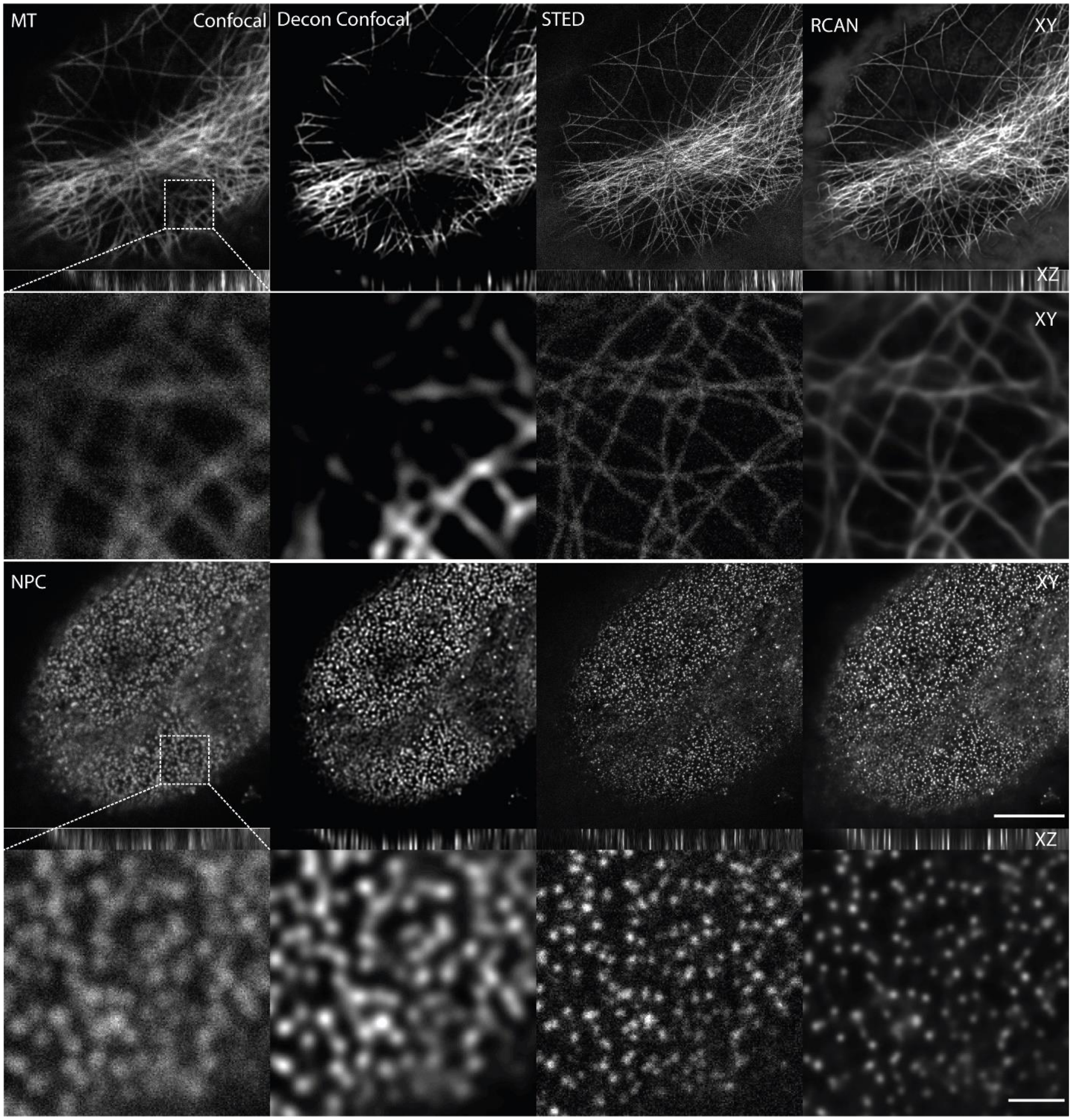


**Supplementary Fig. 12, RCAN provides sharper reconstructions than confocal deconvolution.** Results are shown for fixed microtubules (MT, top) and nuclear pore complexes (NPC, bottom), including lateral (XY) and axial (XZ) slices through image volumes. Second and fourth rows show higher magnification views of rectangular regions in first and third rows. Scale bars: 5 μm lower magnification views and 1 μm higher magnification views.

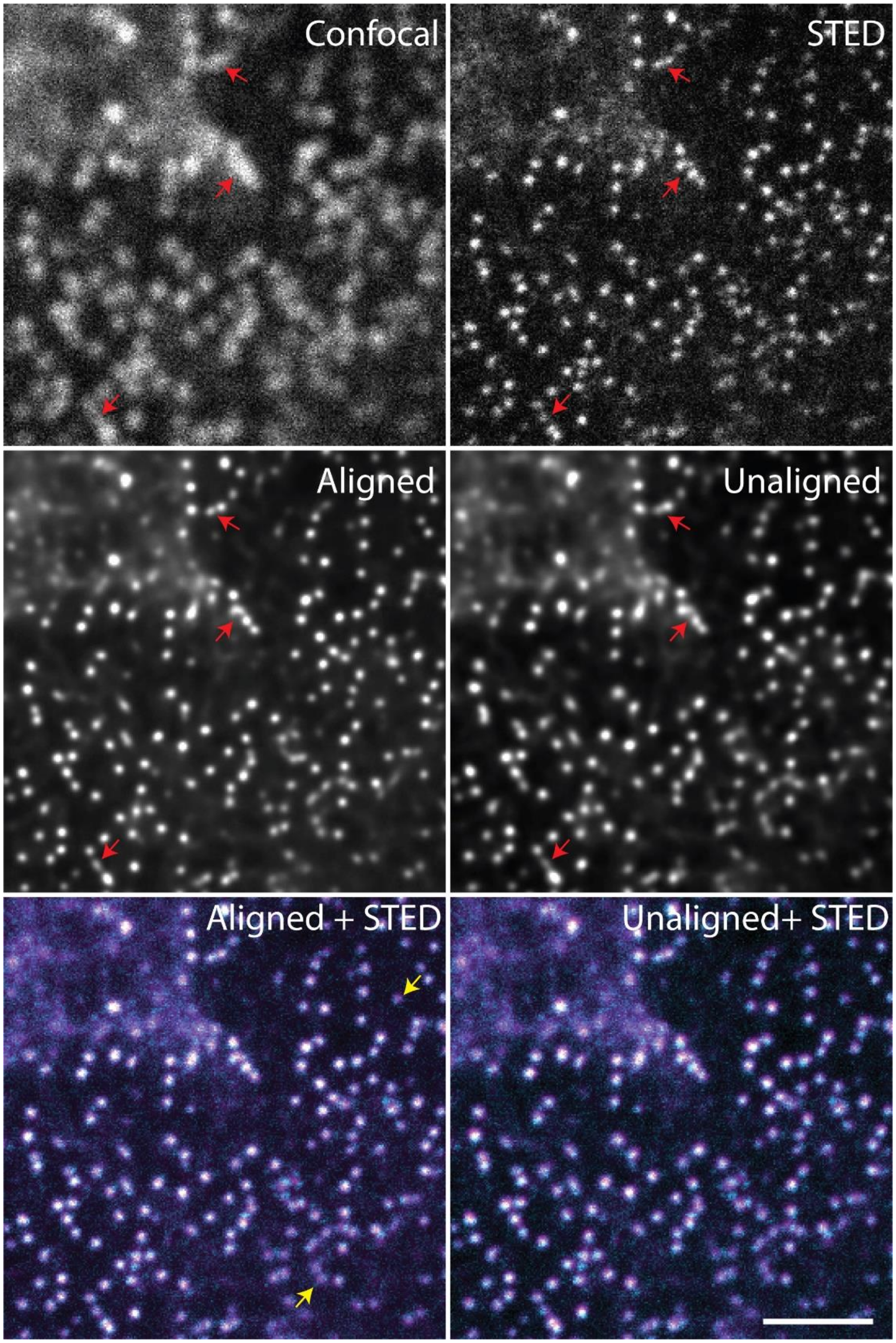


**Supplementary Fig. 13, Alignment improves RCAN prediction.** *Top row:* example confocal (left) and STED (right) microscopy images of nuclear pore complexes. *Middle row*: RCAN predictions when training pairs are aligned (left) or not (right) using an affine registration. Red arrows highlight areas obviously improved after alignment. *Bottom*: cyan/magenta STED/RCAN alignment, showing positional discrepancies (yellow arrows) even after affine registration. Scale bar: 1 μm.

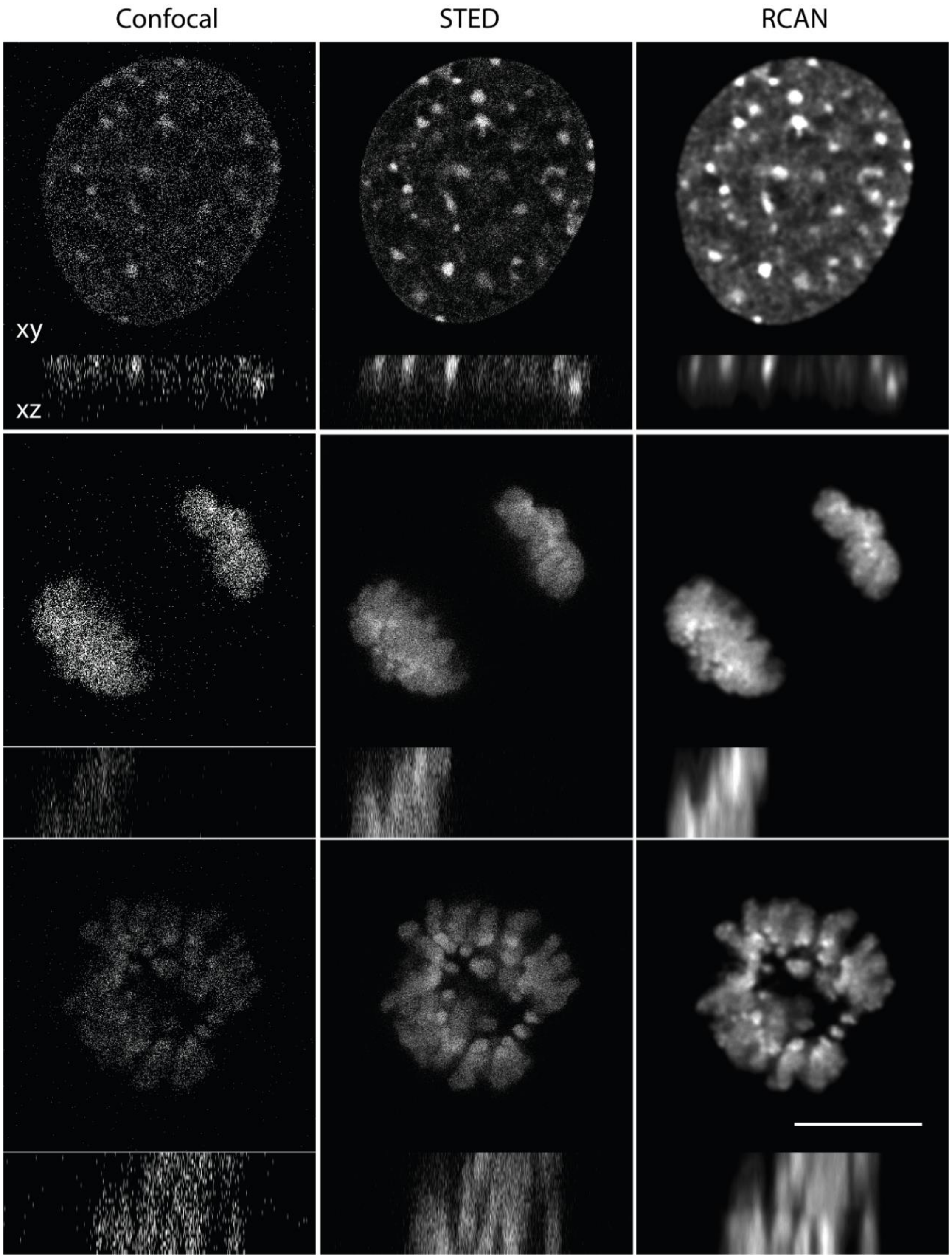


**Supplementary Fig. 14, Additional nuclear morphologies in fixed cells.** Mouse embryonic fibroblasts were fixed stained with SiR-DNA. Rows show different nuclear states with lateral (XY) and axial (XZ) slices. Columns show confocal microscopy images (left), STED microscopy images (middle) and RCAN predictions baed on confocal input (right). Scale bar: 5 μm.

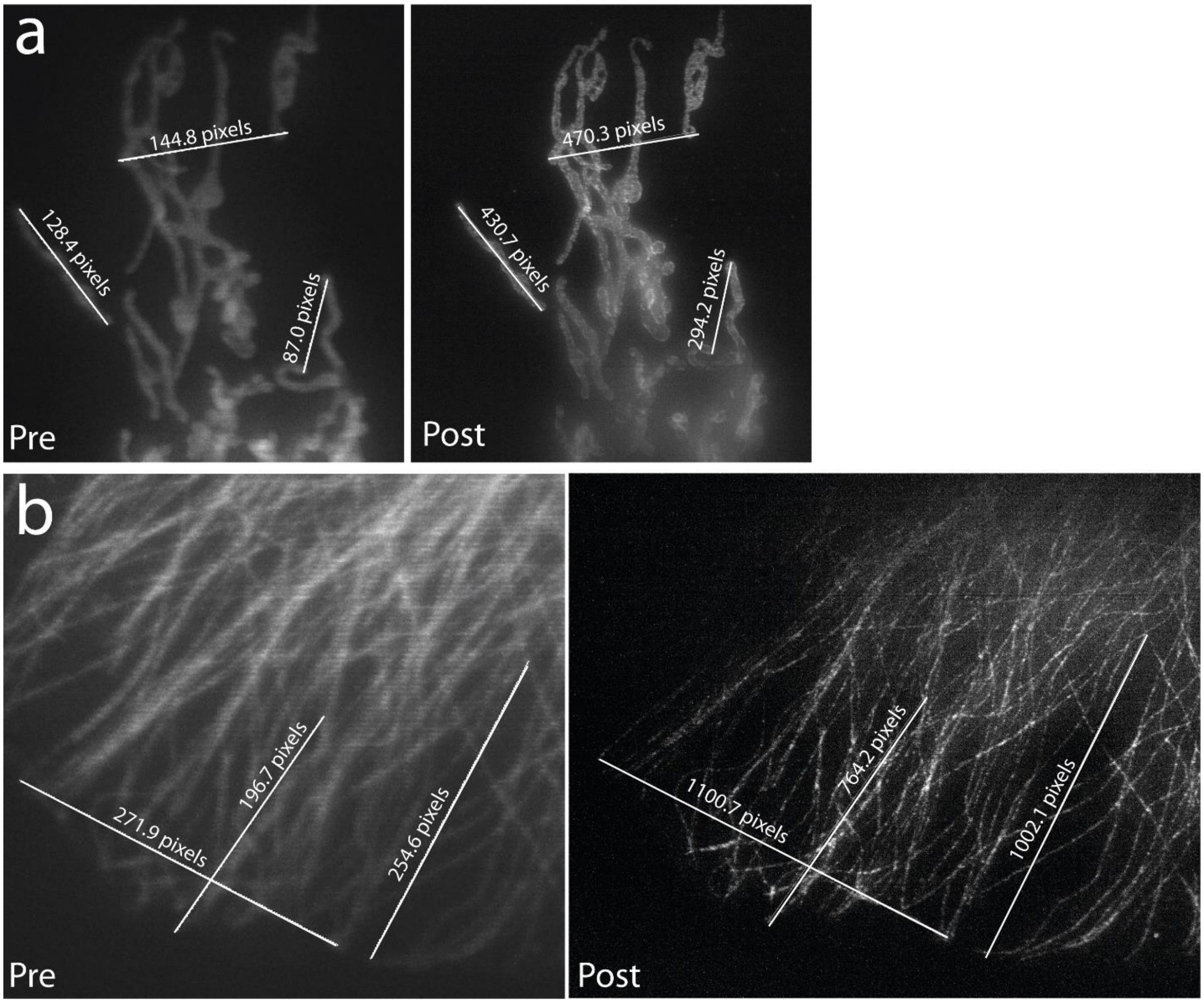


**Supplementary Fig. 15, Example pre- and post-expansion microscopy, imaged on the instant SIM. a)** Mitochondrial label, pre- (left) and post- (right) expansion. **b)** Microtubule label, pre- (left) and post- (right) expansion. Pixel measurements used to estimate expansion factor, for 3 example structures, are shown. Raw (pre-deconvolution) maximum intensity images are shown.

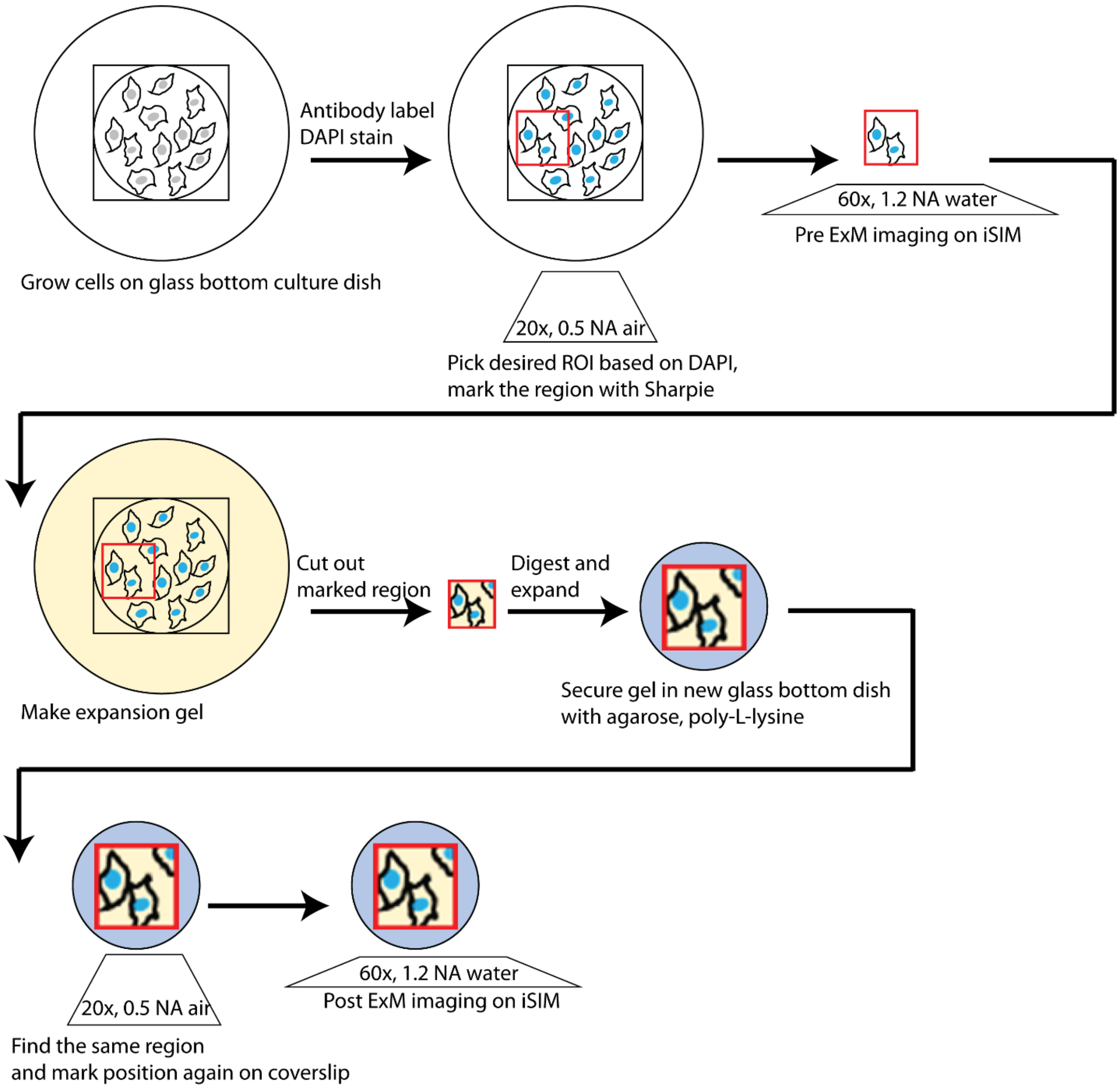


**Supplementary Fig. 16, Pre- and post- ExM workflow.** Cells are grown on glass bottomed dishes, fixed, and labeled with antibody:dye conjugates and DAPI. The labeled samples are imaged at low magnification (0.5 NA, 20x air objective) on a widefield microscope, the desired region of interest (ROI) recorded, and the ROI marked on the bottom of the coverslip with Sharpie marker. This region is then imaged at high NA (1.2 NA, 60x water objective) using the iSIM system. Next, the sample is infiltrated with expansion gel, the marked region cut out, digested, and expanded and secured in a new glass bottom dish with poly-L-lysine and agarose. The ROI of interest is found again under low magnification by comparing the sample to the previously recorded image, the new glass bottom dish marked again with Sharpie, and finally imaged again with iSIM at high NA. See **Methods** for more information.

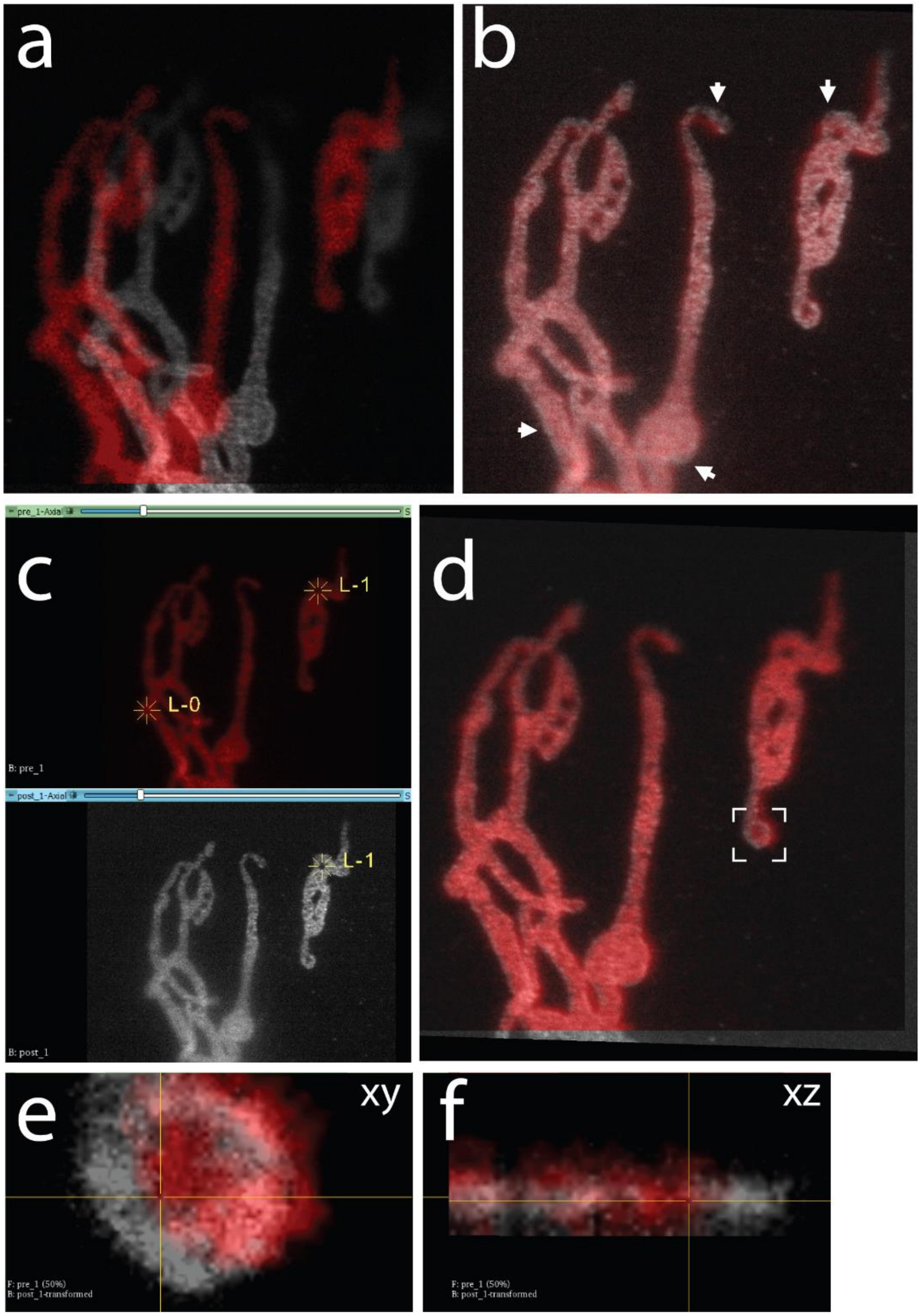


**Supplementary Fig. 17, Attempting to finely register pre- and post- expansion data. a)** Pre-expansion (red) data is scaled up 3.2-fold to coarsely match the post-expansion (grey) data. **b)** Using a 6 degree of freedom rigid affine registration in 3DSlicer results in a coarse registration, with multiple regions (white arrows) that are not registered well. Manually adding landmarks (marked as L-0 and L-1 in **c**) helps to improve registration using a thin plate spline (**d**). However, close inspection of lateral (**e**) and axial (**f**) views at higher magnification (white dashed in box in **d**) reveals that considerable misalignment persists at small length scales. Data are pre-deconvolution images of mitochondria immunolabeled against Tomm20, imaged with iSIM pre- and post- expansion. Screenshots of single slices viewed in 3DSlicer are shown.

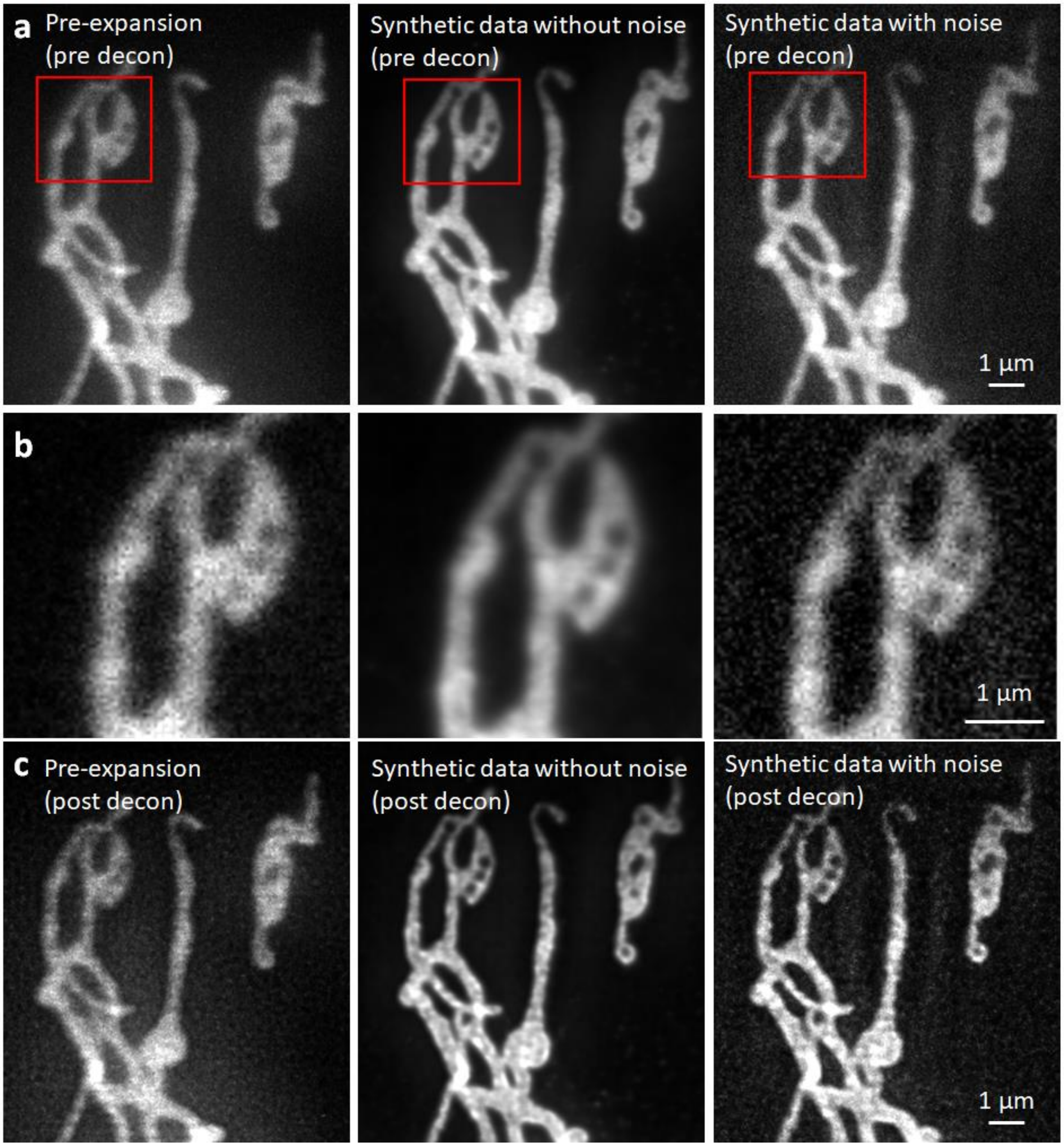


**Supplementary Fig. 18, The effect of adding noise to synthetic data. a)** Pre-expansion data (mitochondria immunolabeled against Tomm20 in fixed U2OS cells acquired on iSIM), synthetic data, and synthetic data with noise added. All data are shown before deconvolution. **b)** Higher magnification views of red boxed region in **a**). **c)** As in **a),** but after deconvolution. See also **Supplementary Fig. 19.**

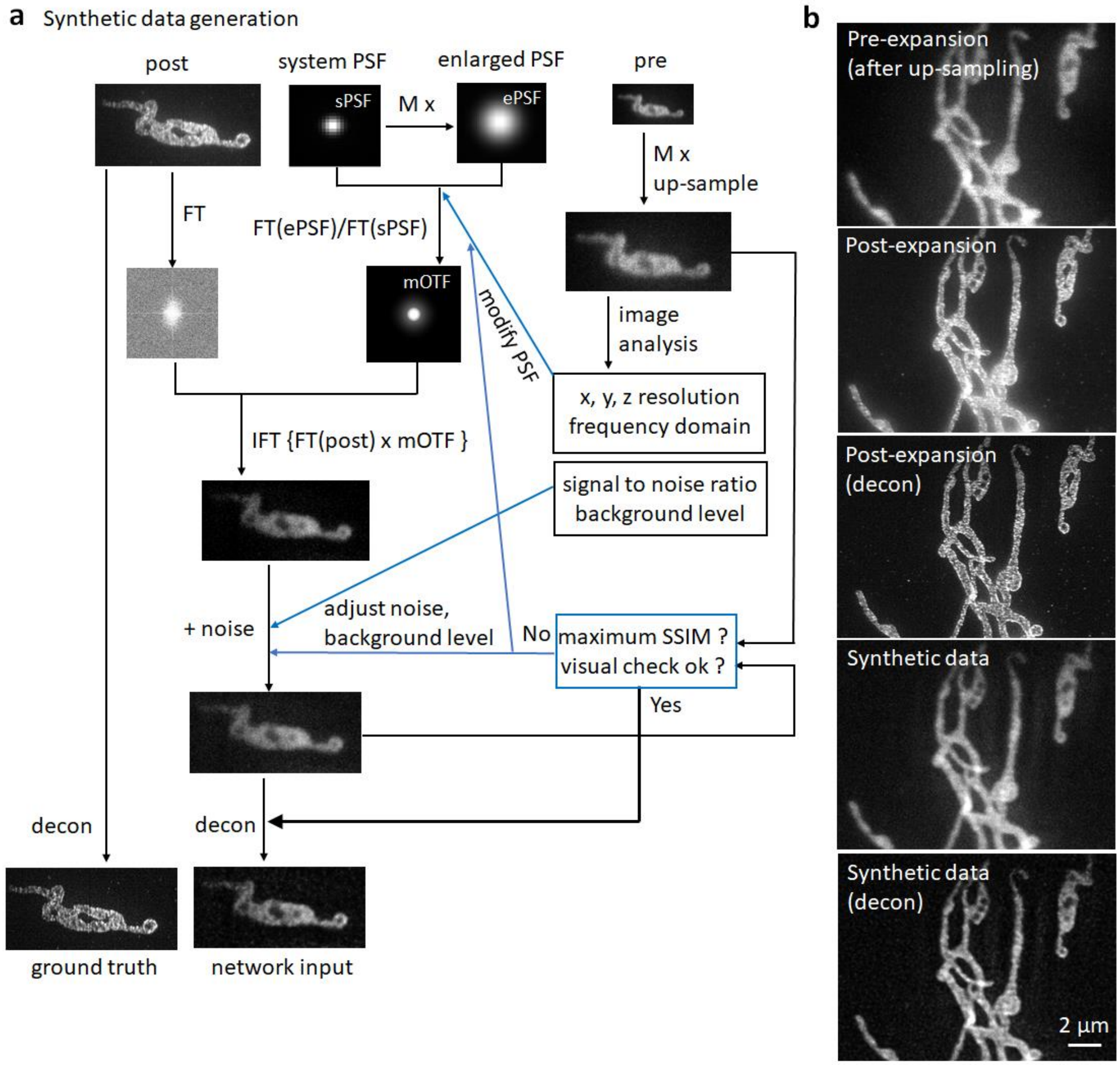


**Supplementary Fig. 19, Generating synthetic pre-expansion data. a)** Schematic workflow for generating synthetic data, used in training RCAN for resolution enhancement. Post-expansion data are blurred, then background and noise are adjusted based on SSIM and visual comparison so that synthetic data resemble acquired pre-expansion data. post: post-expansion data; pre: pre-expansion data; M: expansion factor; sPSF: system PSF; ePSF: enlarged/upsampled PSF; mOTF: modified OTF, i.e., the ratio of FT(ePSF) to FT(sPSF); SSIM: structural similarity index; FT: Fourier transform; IFT: inverse Fourier transform; decon: deconvolution. See **Methods** for more detail. **b)** Comparison between pre-expansion image (after 3.2x upsampling), 3.2x expanded image (before and after deconvolution), and synthetic image (before and after deconvolution) of mitochondria immunolabeled against Tomm20.

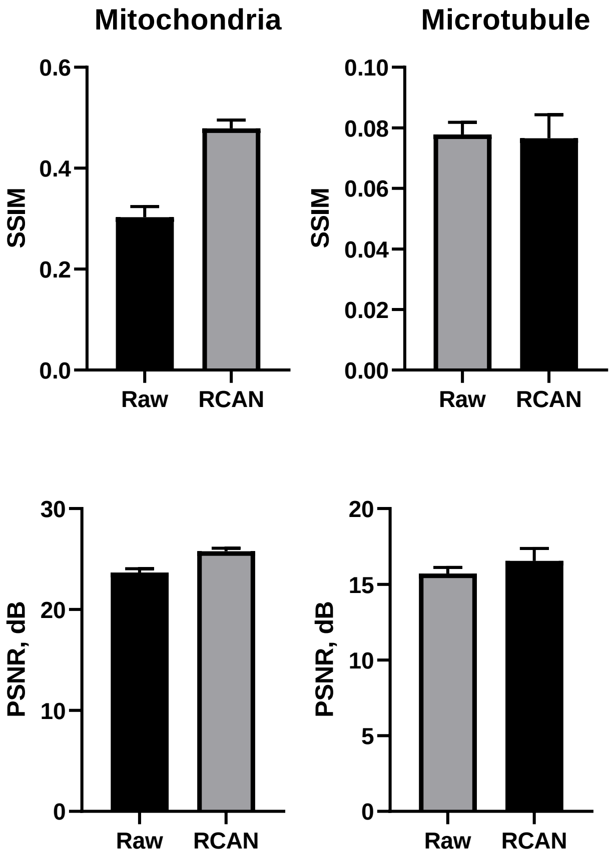


**Supplementary Fig. 20, SSIM and PSNR analysis on synthetic expansion data.** Structural similarity index (SSIM) and peak signal-to-noise ratio (PSNR) for mitochondrial and expanded datasets shown in **Fig. 4b**. Here 'Raw' refers to the synthetic deconvolved iSIM input, and 'RCAN' is the prediction based on this input. See also **Supplementary Table 4**.

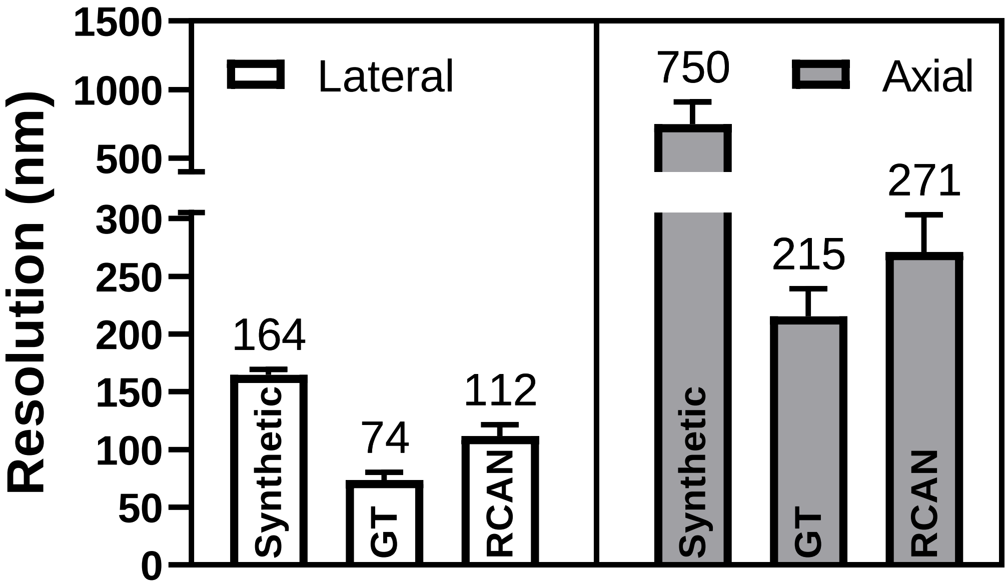


**Supplementary Fig. 21, Resolution analysis for synthetic deconvolved microtubule data.** Average resolution quantification from decorrelation analysis on microtubule samples. Lateral (left) and axial (right) values are shown for synthetic deconvolved iSIM (left columns, 164 +/- 5 nm and 750 +/- 161 nm), ground truth expanded data (middle columns, 74 +/- 6 nm and 215 +/- 24 nm), and RCAN predictions (right columns, 112 +/- 10 nm and 271 +/- 32 nm). Note discontinuous representation of ordinate axis. Mean (shown also above each column) +/- standard deviations derived from 18 (lateral) and 11 (axial) measurements are shown. See also **Fig. 4c**.

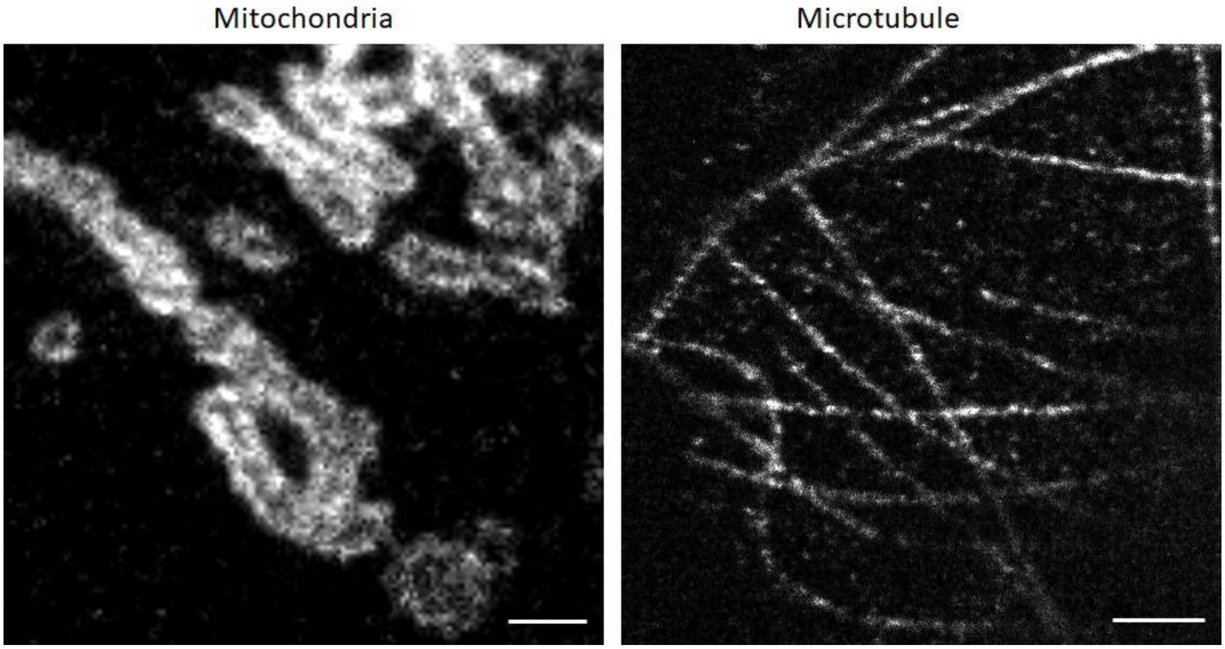


**Supplementary Fig. 22, Variations in label density in expanded samples.** Left: expanded samples immunostained against Tomm20 marking mitochondria; Right: expanded samples immunostained against microtubules. Images are deconvolved single slices. Scale bar: 1 μm.

**Supplementary Table 1, Antibodies and plasmids used in this work.**

| | Figure | Reagent Name | Source | Catalog Number |
|---|---|---|---|---|
| Antibody & Dye | Fig. 4b, Fig. S15a, Fig. S17, Fig. S18, Fig. S19, Fig. S22 | Rabbit-α-Tomm20 | Abcam | ab78547 |
| | | Donkey-α-Rabbit Biotin | Jackson ImmunoResearch | 711-067-003 |
| | | Alexa Fluor 488 Streptavidin | Jackson ImmunoResearch | 016-540-084 |
| | Fig. 4b, Fig. 4e, Fig. S2, Fig. S15b, Fig. S22 | Mouse-α-Alpha Tubulin | Life Technologies | 322500 |
| | | Donkey-α-Mouse Biotin | Jackson ImmunoResearch | 715-065-150 |
| | | Alexa Fluor 488 Streptavidin | Jackson ImmunoResearch | 016-540-084 |
| | Fig. 3a, Fig. 3b, Fig. S12, Fig. S13 | Mouse-α-Nuclear Pore Complex Proteins | Abcam | ab24609 |
| | | Goat-α-Mouse Alexa-594 | ThermoFisher Scientific | A11005 |
| | | Rabbit-α-Alpha Tubulin | Abcam | ab18251 |
| | | Goat-α-Rabbit ATTO-647N | Sigma-Aldrich | 40839 |
| | Fig. S2 | Alexa Fluor 488 Phalloidin | ThermoFisher Scientific | A12379 |
| | Fig. 3c, Fig. 3e, Fig. 3f, Fig. S14 | SiR-DNA | Spirochrome | SC007 |

| | | | | |
|---|---|---|---|---|
| Plasmid | Fig. S2, Fig. S4 | ERmoxGFP | Addgene | 68072 |
| | Fig. S2 | LAMP1-EGFP | Gift from J. Taraska | N/A |
| | Fig. S2 | GalT-GFP | Gift from G. Patterson | N/A |
| | Fig. 4e | EMTB-3XGFP | Addgene | 26741 |
| | Fig. 1f | mApple-Lysosomes-20 (LAMP-1 mApple) | Addgene | 54921 |
| | Fig. 1e, Fig. 1f, Fig. S6 | pShooter pEF-Myc-mito-GFP | Gift from Panagiotis Chandris | N/A |
| | Fig. 1b, Fig. 1c, Fig. 4d, Fig. S1, Fig. S6, Fig. S7 | mEmerald-Tomm20-C-10 | Addgene | 54281 |

**Supplementary Table 2, Acquisition/training parameters for data shown in this work**.

| | Organelle | Modality | Data | | Laser Power | | Z step (µm) | Figure |
|---|---|---|---|---|---|---|---|---|
| | | | Training | Test | Low SNR | High SNR | | |
| Denoising | Actin | iSIM | 20 3D pairs | 5 volumes | 0.3 mW | 33 mW | 0.25 | Fig. S2 |
| | ER | iSIM | 16 3D pairs | 6 volumes | 0.3 mW | 33 mW | 0.25 | Fig. S2, S4 |
| | Golgi | iSIM | 24 3D pairs | 6 volumes | 0.3 mW | 33 mW | 0.25 | Fig. S2 |
| | Lysosome | iSIM | 23 3D pairs | 6 volumes | 0.3 mW | 33 mW | 0.25 | Fig. 1f, Fig. S2 |
| | Microtubule | iSIM | 15 3D pairs | 6 volumes | 0.3 mW | 33 mW | 0.25 | Fig. S2 |
| | Tomm20 Mito | iSIM | 30 3D pairs | 5volumes | 0.3 mW | 33 mW | 0.25 | Fig. 1 b,c, Fig. S1, S2 |
| | Matrix Mito | iSIM | 36 3D pairs | 2600 volumes | 0.3 mW | 33 mW | | Fig. 1e, f |
| Synthetic Resolution Enhancement | Phantom (Synthetic) | N.A. | 23 3D pairs | 7 volumes | N.A. | N.A. | N.A. | Fig. 2 b,c |
| | | | | | Confocal | STED | | |
| Confocal to STED | Microtubule | Confocal & STED | 26 3D pairs | 6 volumes | 10 µW | 40 µW Excitation/ 105 mW Depletion | 0.16 | Fig. 3 a,b |
| | NPC | Confocal & STED | 26 3D pairs | 6 volumes | 16 µW | 23 µW Excitation/ 85 mW Depletion | 0.16 | Fig. 3 a,b |
| | SiR-DNA | Confocal & STED | 22 3D pairs | 11 volumes | 33 µW Live/ 0.5 µW Fixed | 6 µW Excitation/ 35 mW Depletion | 0.16 | Fig. 3 c,e, f |
| | | | | | Pre | Post | | |

| | | | | | | | | |
|---|---|---|---|---|---|---|---|---|
| iSIM to expansion | Tomm20 Mito | iSIM & Expansion | 28 3D pairs | 20 volumes | 9.3 mW | 33 mW | 0.25 | Fig 4, Fig S22 |
| | Microtubule | iSIM & Expansion | 18 3D pairs | 10 volumes | 9.3 mW | 33 mW | 0.25 | Fig 4, Fig S22 |

**Supplementary Table 3, Network comparisons for denoising different organelles.** std: standard deviation. PSNR values are reported in dB.

| | | Networks | | | | | | | | | | | | | | | Figure |
|---|---|---|---|---|---|---|---|---|---|---|---|---|---|---|---|---|---|
| | | **Raw** | | | **CARE** | | | **SRResNet** | | | **ESRGAN** | | | **RCAN** | | | |
| | | Mean | std | N | Mean | std | N | Mean | std | N | Mean | std | N | Mean | std | N | |
| **Actin** | SSIM | 0.37 | 0.02 | 21 | 0.87 | 0.01 | 21 | 0.84 | 0.01 | 21 | 0.78 | 0.01 | 21 | 0.86 | 0.01 | 21 | S3 |
| | PSNR | 25.27 | 0.45 | 21 | 33.56 | 0.83 | 21 | 30.71 | 1.12 | 21 | 29.43 | 0.57 | 21 | 33.60 | 0.59 | 21 | |
| **ER** | SSIM | 0.07 | 0.02 | 11 | 0.52 | 0.02 | 11 | 0.75 | 0.03 | 11 | 0.67 | 0.06 | 11.00 | 0.73 | 0.02 | 11 | S3 |
| | PSNR | 17.22 | 1.94 | 11 | 27.10 | 0.52 | 11 | 26.23 | 1.24 | 11 | 25.88 | 1.38 | 11 | 29.83 | 1.00 | 11 | |
| **Golgi** | SSIM | 0.02 | 0.00 | 14 | 0.41 | 0.03 | 14 | 0.72 | 0.03 | 14 | 0.71 | 0.04 | 14 | 0.74 | 0.01 | 14 | S3 |
| | PSNR | 9.13 | 0.52 | 14 | 19.97 | 1.13 | 14 | 28.59 | 2.42 | 14 | 28.73 | 2.46 | 14 | 32.11 | 1.15 | 14 | |
| **Lysosome** | SSIM | 0.11 | 0.01 | 17 | 0.72 | 0.01 | 17 | 0.83 | 0.01 | 17 | 0.82 | 0.03 | 17 | 0.85 | 0.02 | 17 | S3 |
| | PSNR | 18.28 | 0.58 | 17 | 28.89 | 0.18 | 17 | 33.43 | 0.85 | 17 | 32.36 | 0.98 | 17 | 35.48 | 0.73 | 17 | |
| **MT** | SSIM | 0.12 | 0.03 | 10 | 0.75 | 0.02 | 10 | 0.84 | 0.02 | 10 | 0.78 | 0.02 | 10 | 0.83 | 0.03 | 10 | S3 |
| | PSNR | 17.66 | 1.09 | 10 | 28.25 | 1.73 | 10 | 29.74 | 1.44 | 10 | 27.04 | 1.05 | 10 | 29.85 | 1.73 | 10 | |
| **Tomm20 Mitochondria** | SSIM | 0.15 | 0.03 | 7 | 0.80 | 0.03 | 7 | 0.79 | 0.02 | 7 | 0.80 | 0.02 | 7 | 0.86 | 0.02 | 7 | 1b |
| | PSNR | 17.61 | 0.91 | 7 | 25.30 | 1.03 | 7 | 25.15 | 0.71 | 7 | 25.97 | 0.85 | 7 | 26.75 | 0.87 | 7 | |

**Supplementary Table 4,** SSIM and PSNR values for resolution enhancement. Data for spherical phantoms (**Fig. 2**), confocal to STED (**Fig. 3**), and expansion (**Fig. 4**) are shown. std: standard deviation. PSNR values are reported in dB.

| | | | **2x Deblur** | | | **3x Deblur** | | | **Figure** |
|---|---|---|---|---|---|---|---|---|---|
| | | | Mean | std | N | Mean | std | N | |
| Phantoms | **CARE** | SSIM | 0.94 | 0.004 | 8 | 0.82 | 0.006 | 8 | Fig.2d |
| | | PSNR | 31 | 0.38 | 8 | 23 | 0.26 | 8 | |
| | **SRResNet** | SSIM | 0.93 | 0.003 | 8 | 0.82 | 0.009 | 8 | |
| | | PSNR | 28 | 0.39 | 8 | 23 | 0.29 | 8 | |
| | **ESRGAN** | SSIM | 0.91 | 0.002 | 8 | 0.81 | 0.003 | 8 | |
| | | PSNR | 28 | 0.31 | 8 | 23 | 0.22 | 8 | |
| | **RCAN** | SSIM | 0.98 | 0.002 | 8 | 0.93 | 0.004 | 8 | |
| | | PSNR | 38 | 0.24 | 8 | 32 | 0.36 | 8 | |
| | **Raw** | SSIM | 0.65 | 0.008 | 8 | 0.46 | 0.011 | 8 | |
| | | PSNR | 19 | 0.35 | 8 | 14.16 | 0.35 | 8 | |
| | | | **MT** | | **NPC** | | **SiR-DNA** | | |
| | | | Confocal | RCAN | Confocal | RCAN | Confocal | RCAN | |
| Confocal To STED | **SSIM** | Mean | 0.19 | 0.33 | 0.22 | 0.39 | 0.52 | 0.63 | Fig.S10 |
| | | Std | 0.04 | 0.03 | 0.03 | 0.03 | 0.11 | 0.09 | |
| | | N | 20 | 20 | 22 | 22 | 33 | 33 | |
| | **PSNR** | Mean | 16 | 19 | 16 | 21 | 18 | 24 | |
| | | Std | 1 | 1 | 1 | 1 | 1 | 2 | |
| | | N | 20 | 20 | 22 | 22 | 33 | 33 | |
| | | | **MT** | | **Mitochondria** | | | | |
| Expansion | **SSIM** | | Raw | RCAN | Raw | RCAN | | | Fig. S20 |
| | | Mean | 0.08 | 0.08 | 0.30 | 0.48 | | | |
| | | Std | 0.00 | 0.01 | 0.02 | 0.02 | | | |
| | | N | 19 | 19 | 14 | 14 | | | |
| | **PSNR** | Mean | 16 | 17 | 24 | 26 | | | |
| | | Std | 0.4 | 0.8 | 0.4 | 0.3 | | | |
| | | N | 19 | 19 | 14 | 14 | | | |

**Supplementary Note 1, Image Restoration and Enhancement Using 3D Residual Channel Attention Networks**

### 1. Image Restoration and Enhancement Using Deep Neural Networks

Image restoration generally attempts to find a mapping $\mathcal{F}(\mathbf{y}) = \mathbf{x}$ that recovers a latent clean image $\mathbf{x}$ from a degraded observation $\mathbf{y}$, assuming an image degradation model

$$\mathbf{y} = \mathbf{x} + \mathbf{v}$$

where $\mathbf{v}$ is a residual component representing image degradation such as noise and blurring. Zhang *et al*.[1] proposed an image denoising method based on residual learning where a residual mapping $\mathcal{R}(\mathbf{y}) = \mathbf{v}$ is learnt instead of the original mapping $\mathcal{F}(\mathbf{y})$, using a deep convolutional neural network with a long skip connection. The residual mapping is generally easier to optimize than the original identity mapping[2] and often allows deeper networks with skip connections to be used.

The problem of image enhancement such as reconstructing an accurate high-resolution (HR) image given its low-resolution (LR) counterpart is referred to as single image super-resolution (SR)[3]. Such image SR is an ill-posed inverse problem since multiple HR solutions exist for a given LR input. To tackle this problem, numerous deep convolutional neural network (CNN) based methods have been proposed to learn mappings between LR and HR image pairs. Pioneering work in this area was performed by Dong et al.[4], who proposed SRCNN for image SR using a three-layer CNN. Ledig *et al.*[5] utilized the residual learning approach for single image super-resolution with deep residual network and skip connection (SRResNet). Lim *et al*.[6] refined SRResNet by employing a better and simplified residual block structure and proposed an image super-resolution network named enhanced deep super-resolution (EDSR).

In single image SR, the goal is to retrieve as much high-frequency information as possible from its LR input. However, the LR input images are dominated by low spatial frequency information, which can be directly forwarded to the final HR outputs as they do not require much additional processing. High-frequency channel-wise features are more informative for HR enhancement, but most CNN-based methods treat channel-wise features equally, which leads to low resolution features dominating the CNN output. Attention mechanisms can bias the allocation of available processing resources towards the most informative components of an input[7]. EDSR was further improved by Zhang *et al*.[8] by incorporating the channel attention module in each residual block named residual channel attention networks (RCAN). Although RCAN was originally designed for single image super-resolution, we found that it is also very effective for image restoration tasks such as denoising.

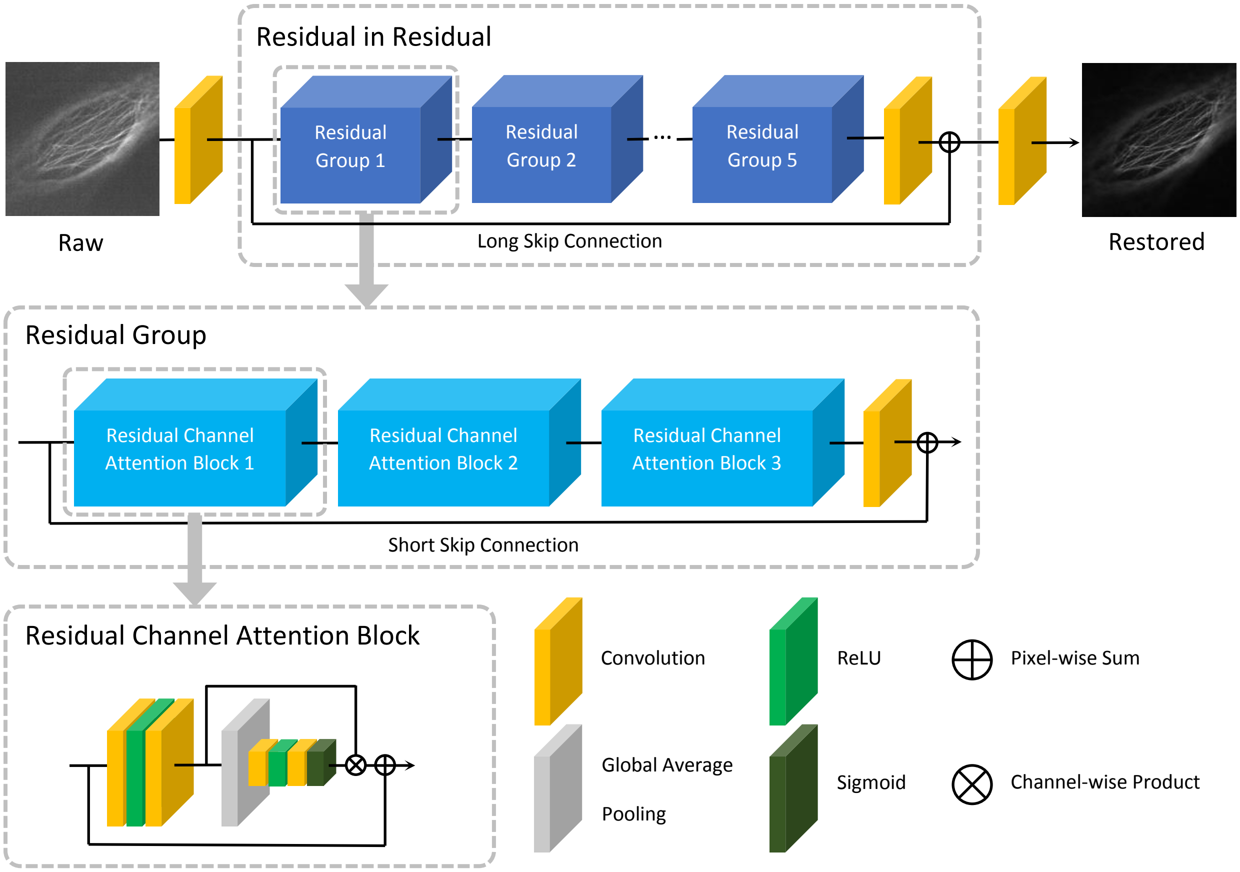


**Fig. 1. 1. Network architecture of the 3D residual channel attention network (3D RCAN).** ReLU: Rectified linear unit. See text for further information, see also **Fig. 1a**.

### 2. 3D RCAN architecture

We extended the original RCAN to handle image volumes, using smaller numbers of residual groups (RG), residual channel attention blocks (RCAB) and convolution layers than the original RCAN implementation[8] so that the model fits into GPU memory. The overall network architecture of 3D RCAN is the same as the original RCAN except that we omitted the upscaling module at the end of the network so that the output of the model has the same size as the input. The network architecture of the 3D RCAN is shown in **Fig. 1.1**.

The 3D RCAN consists of three modules which perform shallow (single layer) feature extraction, residual in residual (RIR) feature extraction, and reconstruction. First, the shallow feature $F_{SF}$ is extracted from the input image $I_{in}$ as

$$F_{SF} = H_{SF}(I_{in})$$

where $H_{SF}$ denotes the convolution before the RIR module (in **Fig. 1.1** convolutional layer immediately after input layer). Next, deep features $F_{DF}$ are extracted by the RIR module as

$$F_{DF} = H_{RIR}(F_{SF}).$$

Finally, the output image $I_{out}$ is reconstructed by the convolution $H_{REC}$ at the end of the network (in **Fig. 1.1** convolutional layer immediately prior to output) as

$$I_{out} = H_{REC}(F_{DF}).$$

The RIR operation is formulated as

$$H_{RIR}(F_{SF}) = F_{SF} + H\left(H_{RG}^{5}\left(H_{RG}^{4}\left(\ldots\ H_{RG}^{1}(F_{SF})\right)\right)\right)$$

where $H_{RG}^{i}$ is the $i$-th RG and $H$ is the convolution after the last RG. The $i$-th RG operation is defined as

$$H_{RG}^{i}\left(F^{i-1}\right) = F^{i-1} + H^{i}\left(H_{RCAB}^{i,3}\left(H_{RCAB}^{i,2}\left(H_{RCAB}^{i,1}\left(F^{i-1}\right)\right)\right)\right)$$

where $F^{i-1}$, $H^{i}$, and $H_{RCAB}^{i,j}$ are the output feature from the $i-1$-th RG, the convolution after the last RCAB within the $i$-th RG, and the $j$-th RCAB within the $i$-th RG, respectively. The $j$-th RCAB operation within the $i$-th RG is defined as

$$H_{RCAB}^{i,j}\left(F^{i,j-1}\right) = F^{i,j-1} + H_{CA}^{i,j}\left(H^{i,j,2}\left(\delta\left(H^{i,j,1}\left(F^{i,j-1}\right)\right)\right)\right)$$

where $F^{i,j-1}$ is the output feature from the previous RCAB, $H^{i,j,1}$ and $H^{i,j,2}$ are two convolutions in this RCAB, $\delta$ is the rectified linear unit (ReLU), and $H_{CA}^{i,j}$ is the channel attention operation. Lastly, the channel attention operation is defined as

$$H_{CA}^{i,j}(F) = F \cdot \sigma\left(H_{U}^{i,j}\left(\delta\left(H_{D}^{i,j}(H_{GP}(F))\right)\right)\right)$$

where $F$ is the input feature to this operation, $H_{GP}$ is the global average pooling, $H_{D}^{i,j}$ and $H_{U}^{i,j}$ are convolutions with channel downscaling and upscaling, and $\sigma$ is the sigmoid gating function.

### 3. Preprocessing of training images

#### 3.1 Image normalization

The dynamic range of raw and ground truth images are typically very different. To normalize images to the same intensity range, we used a percentile-based normalization method[9]. The normalization function for an image $I$ is defined as

$$N\left(I; p_{low}, p_{high}\right) = \frac{I - \mathrm{perc}(I, p_{low})}{\mathrm{perc}\left(I, p_{high}\right) - \mathrm{perc}(I, p_{low}) + \epsilon}$$

where $\mathrm{perc}(I, p)$ is the $p$-th percentile of all pixel values in $I$ and $\epsilon$ is a small scalar value to avoid zero-division. We set $p_{low} = 2, p_{high} = 99.9$, and $\epsilon = 10^{-6}$ in all experiments.

#### 3.2 Image alignment

We sometimes observed small displacements between the raw and ground truth images. To train deep learning models for image restoration (and supervised image-to-image translation in general), it is important that raw and ground truth images are precisely aligned. We implemented a 3D version of the phase correlation algorithm[10] and used it for image alignment prior to training.

### 4. Implementation details and training settings

We implemented a 3D version of RCAN[8] using Keras[11] with a TensorFlow[12] backend. During the training process, a pair of raw and ground truth patches with size $256 \times 256 \times 16$ is randomly cropped from training data, then augmented by random rotation and flipping. The patch size used in our implementation is much larger than the one used in the original RCAN ($48 \times 48$). We observed that smaller patches led to unstable training and poor restoration results. We suspect that this is because microscopy images may show less high spatial frequency content than natural images. A larger patch is necessary to extract enough gradient information for back-propagation.

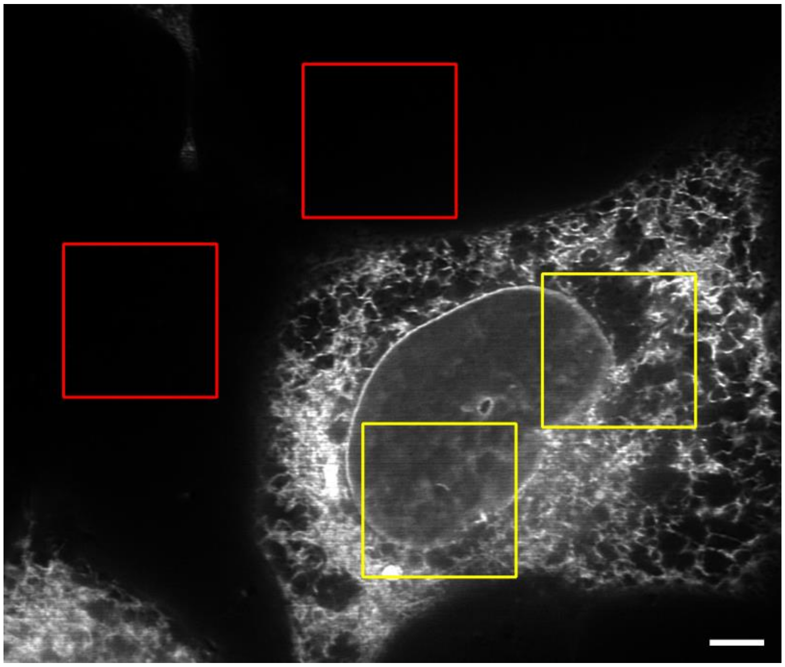

**Fig. 1.2, Background patch rejection.** Examples of automatically detected background (red) and foreground (yellow) patches (scale bar: 5 μm).

In microscopy images, foreground objects of interest might be distributed sparsely. In this case, the model might overfit the background area and fail to learn the structure of the foreground objects if the entire image is used indiscriminately for training. To avoid overfitting, patches from the background regions are excluded from training (**Fig. 1.2**). To determine whether a patch pair is from the foreground region, thresholding with a threshold value $\tau$ is applied to a ground truth patch. Pixels with intensity values greater than $\tau$ are considered foreground. Afterwards, the foreground pixel ratio

$$r = \frac{\#\text{foreground pixels in the patch}}{\#\text{total pixels in the patch}}$$

is calculated. A patch pair is considered to come from the foreground regions if and only if $r \geq \rho$. We set $\tau = 0.25$ and $\rho = 0.05$ in all experiments.

The model is optimized by minimizing the $L_1$ loss

$$L_1(\Theta) = \frac{1}{N}\sum_{i=1}^{N}\left|f\left(I_{raw}^{i};\Theta\right) - I_{gt}^{i}\right|$$

where $f$ and $\Theta$ denote the 3D RCAN and its parameters, $I_{raw}^{i}$ and $I_{gt}^{i}$ are the $i$-th pair of raw and ground truth image patches, and $N$ is the total number of patch pairs. The Adam optimizer[13] with $\beta_1 = 0.9, \beta_2 = 0.999,$ and $\epsilon = 10^{-7}$ is used for training. The learning rate is initialized as $10^{-4}$ and halved at every 100 epochs. 256 patch pairs are processed in each epoch. Each model was trained on two NVIDIA GeForce GTX 1080 Ti GPUs for 400 epochs, which took 1 day. Applying the denoising model on a 1920 x 1550 x 12 dataset using a desktop with a single GTX 1080 Ti GPU took 63.3 s per volume. This time includes the time it takes to save the volume (using 32-bit output). On similar datasets with the same XY dimensions (but different number of Z-slices), applying the model took 3.9 s to 5.2 s per Z-slice.

### 5. Model uncertainty estimation

We followed the procedure proposed by Weigert *et al*[9] to estimate the uncertainty in model prediction by using a normalized disagreement score. To compute the disagreement among multiple models, we extended the RCAN model to output a distribution of values for each pixel, instead of just a single value. Specifically, the RCAN model is extended to produce two output channels that represent location $\mu$ and scale $\sigma$ parameters of the Laplace distribution of each pixel

$$p(x;\mu,\sigma) = \frac{1}{2\sigma}\exp\left(-\frac{|x-\mu|}{\sigma}\right).$$

This extended model is optimized by minimizing the average negative log-likelihood.

We independently trained $M$ extended RCAN models[9], where $M = 5$. The mixture distribution $q$ is defined as the average of $M$ predicted distributions

$$q = \frac{1}{M}\sum_{m=1}^{M} p^m$$

where $p^m$ is the Laplace distribution predicted by the $m$-th model. The disagreement score among multiple models is defined as the average Kullback-Leibler divergence from the mixture distribution to each model's distribution

$$\mathcal{D} = \frac{1}{M}\sum_{m=1}^{M} D_{KL}\left(p^m || q\right).$$

Using the fact that $0 \leq \mathcal{D} \leq \log M$, the normalized disagreement score can be defined as

$$\widehat{\mathcal{D}} = \frac{1}{\log M}\mathcal{D} = \frac{1}{M \log M}\sum_{m=1}^{M} D_{KL}\left(p^m || q\right).$$


1 Zhang, K., Zuo, W., Chen, Y., Meng, D. & Zhang, L. Beyond a Gaussian Denoiser: Residual Learning of Deep CNN for Image Denoising. *IEEE Transactions on Image Processing* **26.7**, 3142-3155 (2017).

2 He, K., Zhang, X., Ren, S. & Sun, J. Deep residual learning for image recognition. *IEEE Conference on Computer Vision and Pattern Recognition*, 770-778 (2016).

3 Freeman, W. T., Pasztor, E.C., Carmichael, O.T. Learning Low-Level Vision. *International Journal of Computer Vision* **40**, 25-47 (2000).

4 Dong, C., Loy, C. C., He, K. & Tang, X. Learning a Deep Convolutional Network for Image Super-Resolution. *European Conference on Computer Vision*, 184-199 (2014).

5 Ledig, C. *et al.* Photo-Realistic Single Image Super-Resolution Using a Generative Adversarial Network. *IEEE Conference on Computer Vision and Pattern Recognition*, 4681-4690 (2017).

6 Lim, B., Son, S., Kim, H., Nah, S. & Lee, K. M. Enhanced Deep Residual Networks for Single Image Super-Resolution. *IEEE Conference on Computer Vision and Pattern Recognition Workshops*, 1132-1140 (2017).

7 Hu, J., Shen, L. & Sun, G. Squeeze-and-Excitation Networks. *IEEE/CVF Conference on Computer Vision and Pattern Recognition*, 7132-7141 (2018).

8 Zhang, Y. *et al.* Image Super-Resolution Using Very Deep Residual Channel Attention Networks. *European Conference on Computer Vision*, 286-301 (2018).

9 Weigert, M. *et al.* Content-aware image restoration: pushing the limits of fluorescence microscopy. *Nat Methods* **15**, 1090-1097 (2018).

10 Reddy, B. S. & Chatterji, B. N. An FFT-based technique for translation, rotation, and scale-invariant image registration. *IEEE Trans Image Process.* **5**, 1266-1271 (1996).

11 Chollet, F. & others. *keras*, 2015).

12 Abadi, A. *et al.* TensorFlow: Large-Scale Machine Learning on Heterogenous Distributed Systems. *arXiv*, 1603.04467 (2016).
13 Kingma, D. P. & Ba, J. Adam: A Method for Stochastic Optimization. *ArXiv*, 1412.6980 (2014).
